# *In vivo* CRISPR-based screen identifies ZC3H12C as a mediator of CAR-T cell dysfunction in solid tumors

**DOI:** 10.64898/2026.04.30.721530

**Authors:** Paula Barbao, Alba Rodriguez-Garcia, Ariel Galindo-Albarrán, Marta Giménez-Alejandre, Pablo Clavero, Teresa Lobo-Jarne, Marta Botas, Guillem Colell, Joan Castellsagué, Guim Cascalló, Irene Andreu-Saumell, Marta Soria-Castellano, Salut Colell, Berta Marzal, Beatriz Martín-Mur, Anna Esteve-Codina, Alvaro Urbano-Ispizua, Aleix Prat, Luca Gattinoni, Marco Mendoza-Parra, Sonia Guedan

## Abstract

CAR-T cell therapy has shown limited efficacy in solid tumors, largely due to T cell dysfunction driven by chronic antigen exposure. To uncover mediators of this dysfunction, we developed an *in vivo* screening platform using an ovarian xenograft tumor model in which CD28-based CAR-T cells undergo exhaustion leading to tumor escape. Transcriptomic profiling of tumor-infiltrating CAR-T cells at different stages revealed dynamic upregulation of exhaustion-associated genes. We used this data to design a focused CRISPR/Cas9 library and performed an *in vivo* screen. We identified 14 significantly enriched candidate genes, among which ZC3H12C emerged as the top hit. Single-cell RNA and ATAC-seq confirmed ZC3H12C expression in CAR-T cells undergoing early exhaustion *in vivo*. ZC3H12C disruption enhanced CAR-T cell persistence and antitumor efficacy while reducing exhaustion, across both CD28- and 4-1BB-based CARs targeting distinct antigens. These results highlight ZC3H12C as a promising target to improve CAR-T therapy in solid tumors.

## Introduction

Chimeric antigen receptor (CAR) T cell therapy has demonstrated unprecedented clinical success in patients with hematologic malignancies; however, its efficacy in solid tumors remains limited. Recent reports involving patients with advanced, treatment-refractory solid tumors have shown that CAR-T cells can induce objective antitumor responses^1–8^. Following infusion, CAR-T cells are detectable in peripheral blood, and, in some cases, expand and persist overtime^2^. Infused CAR-T cells in most cases are able to traffic to the tumors^4,6–9^, where they can become activated and proliferate locally in the tumor microenvironment^9^. Despite this, sustained tumor regression and durable clinical responses remain uncommon. While antigen loss has been reported in certain tumor types and targets^4^, in many cases, disease progression occurs despite continued expression of the targeted antigen^1,2,9^. These findings indicate that a key barrier to efficacy in solid tumors is the limited functionality of CAR-T cells.

T cell exhaustion is a differentiation state that has been described in the context of chronic viral infections and cancer. In these settings, persistent antigen stimulation leads to the progressive loss of T cell effector functions, contributing to disease progression^10^. Similar to endogenous T cells, CAR-T cells are also susceptible to exhaustion, which may arise from specific CAR design driving tonic signaling^11^ or from chronic antigen exposure in the tumor site^12^. The mechanisms driving loss of functions in CAR-T cells have been mainly studied in the context of *in vitro* models^13–17^, however, no *in vivo* models have been established that faithfully recapitulate the features of CAR-T cell exhaustion due to chronic antigen exposure in solid tumors.

Here, we set out to dissect the molecular mechanisms driving CAR-T cell dysfunction in solid tumors. We employed an *in vivo* model of solid tumor xenografts in which CAR-T cells infiltrate tumors but progressively lose function, leading to therapy failure despite antigen persistence. Through comprehensive phenotypic, functional and transcriptomic analyses, including bulk and single cell transcriptomic analysis, we characterized the onset and progression of CAR-T cell exhaustion after chronic antigen stimulation in our model. Based on gene expression dynamics in dysfunctional CAR-T cells, we designed a focused CRISPR/Cas9 library screen targeting 300 candidate genes and performed an *in vivo* loss-of-function screen. This approach uncovered 14 previously unrecognized mediators of CAR-T cell dysfunction. We validated three targets, including ZC3H12C, TG, and ITGB8, whose disruption enhanced *in vivo* CAR-T cell persistence and antitumor activity against solid tumors. These findings provide new insights into the mechanisms driving CAR-T cell dysfunction and highlight potential targets to enhance CAR-T therapy efficacy in solid tumors.

## Results

### Profiling CAR-T cell exhaustion induced by chronic antigen exposure in a HER2+ ovarian cancer xenograft model

To study the T cell-intrinsic mechanism that causes CAR-T dysfunction, we used an *in vivo* model of xenograft tumors that recapitulated disease recurrence in absence of antigen loss. NOD-SCID gamma (NSG) mice bearing ovarian subcutaneous tumors (SKOV3) were treated with a single dose of CD28-based CAR-T cells targeting HER2 with low affinity by using a mutated version of the trastuzumab-based 4D5 scFv (4D5.5)^18^. In this model, CAR-T cell therapy induced antitumor responses in all treated mice, but in most cases tumors were not completely eliminated and eventually escaped to therapy (**Fig. 1A**). Post-relapse tumors retained robust HER2 expression and showed marked CD8+ T cell infiltration (**Fig. S1A-B**).

**Figure 1.**
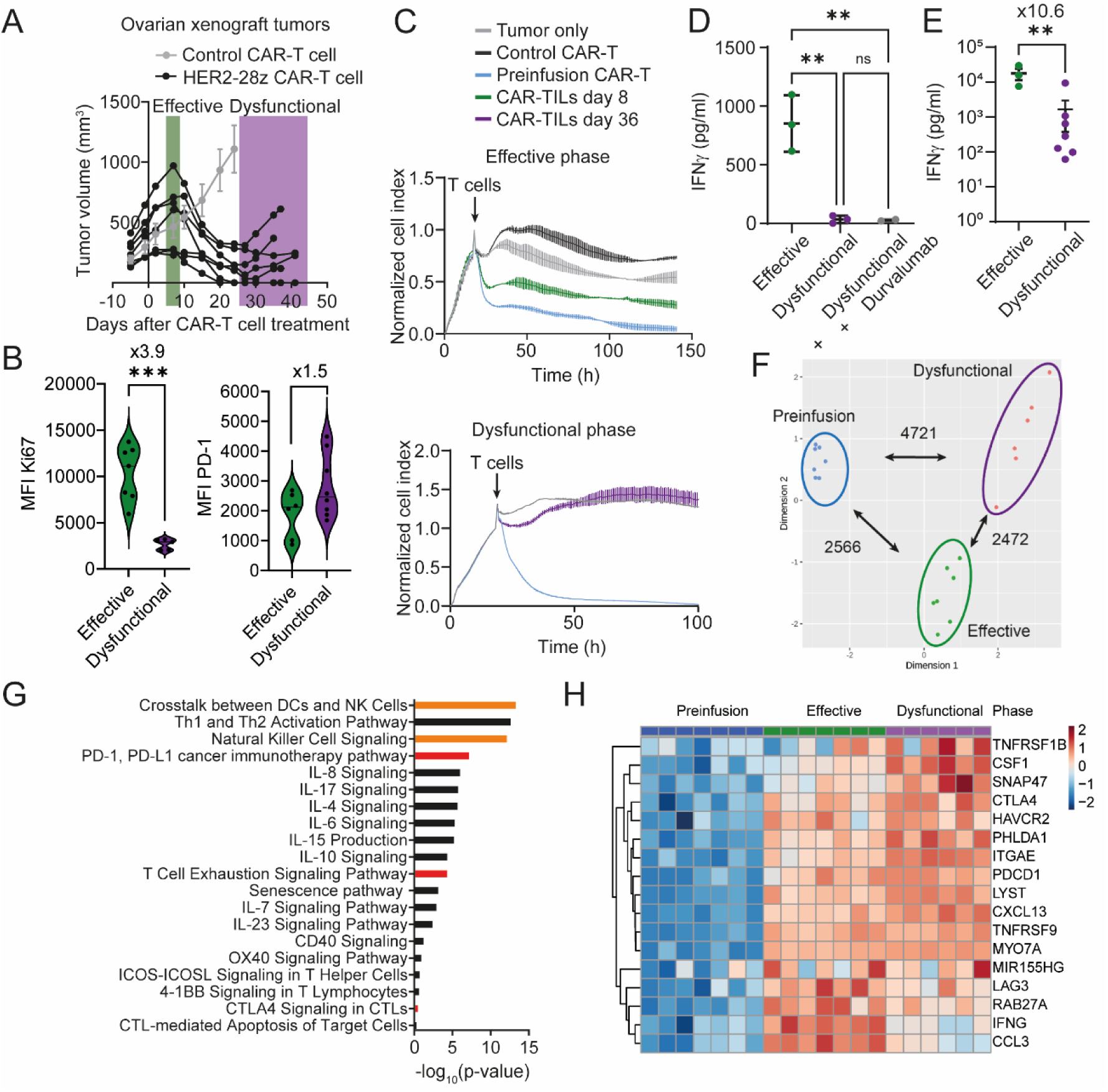
HER2-28z CAR-T cells become dysfunctional after chronic antigen exposure *in vivo* causing tumor escape. NSG mice bearing SKOV3 tumors were treated with a single dose of 2×10^6^ HER2-28z or CD19-28z CAR+ T cells (Control CAR-T cell). **A**) Tumors were measured at indicated time points. Data are plotted as mean ±SEM of tumor volume (control CAR-T, n=6 tumors) or as individual tumors (HER2-CAR-T, n=7 tumors). Representative of n>5 experiments using different healthy donors. Timepoints at which CAR-TILs were isolated from tumors for use in downstream analyses are indicated. **B**) Violin plots showing mean fluorescence intensity of PD-1 (right panel) and Ki67 (left panel) in effective compared to dysfunctional CAR-TILs. Each dot represents a tumor (n=4-5 healthy donors). **C**) *Ex vivo* real-time cytotoxicity analysis of isolated CAR-TILs at effective (upper panel) and dysfunctional (bottom panel) phases against SKOV3 tumor cells (E:T=5:1). Data are plotted as mean ±SEM of normalized cell index of a representative experiment using one healthy donor. Time when T cells were added is indicated. **D**) Analysis of IFNγ production by CAR-TILs after 24h of *ex vivo* co-culture with SKOV3 tumor cells (E:T=3:1). The PD-1/PD-L1 axis was inhibited by the addition of PD-L1 blocking antibody Durvalumab. Data are plotted as mean ±SEM (n=3 in effective, n=2 in dysfunctional and dysfunctional + durvalumab). Each dot represents a tumor used for isolating CAR-TILs. **p<0.01 by one-way ANOVA with Tukey’s multiple comparisons test. **E**) IFNγ secretion by CAR-TILs after overnight stimulation with PMA/Ionomycin. Each dot represents a tumor used for isolating CAR-TILs (n=2 healthy donors). Fold change of effective vs dysfunctional is indicated. **B and E**) **p<0.01, ***p<0.0001 by two-tailed unpaired t-test. Fold change relative to effective (Ki67, IFNγ) or dysfunctional (PD-1) is shown. **F**) Principal component analysis of HER2-28z CAR-T samples from preinfusion, effective, and dysfunctional phases included in the RNA-seq analysis. Number of differentially expressed genes (p-adj. value<0.05 and logFc<1 or logFc<-1) between groups is indicated (n=7 healthy donors for preinfusion; n=6 for effective; n=5 for dysfunctional). **G**) Ingenuity Pathway Analysis (IPA) of upregulated genes in dysfunctional compared to preinfusion samples. Significant selected pathways are ranked by p-value of enrichment. Orange bars denote NK cell-related pathways, and red bars indicate pathways of T cell exhaustion. **H**) Heatmap showing the expression level of 17 TIL marker genes^14^ in preinfusion, effective and dysfunctional samples.

To elucidate whether CAR-T cell dysfunction was responsible for tumor escape in our model, we next isolated CAR tumor infiltrating lymphocytes (CAR-TILs) after infusion (effective phase) and after therapy failure (dysfunctional phase) (**Fig. 1A**). We then analyzed the phenotype and *ex vivo* function of isolated effective and dysfunctional CAR-TILs. We observed higher infiltration of T cells in tumors at dysfunctional compared to effective phase (**Fig. S1D-E**), suggesting initial T cell expansion at the tumor site. However, dysfunctional CAR-TILs showed increased expression of PD-1 and significantly reduced levels of Ki67 (**Fig. 1B and S1C**) compared to effective CAR-TILs. When *ex vivo* co-cultured, effective CAR-TILs efficiently eliminated SKOV3 tumor cells, similar to preinfusion HER2-28z CAR-T cells, whereas dysfunctional CAR-TILs displayed impaired killing capacity (**Fig. 1C**). In line with this, dysfunctional CAR-TILs secreted significantly lower levels of effector cytokines, including IFNγ, compared to effective CAR-TILs in both CAR-dependent (**Fig 1D and S1F**) and CAR-independent settings (**Fig. 1E and S1G**). Blocking the PD-1/PD-L1 axis failed to reinvigorate T cell functions, indicating an advanced exhaustion state (**Fig. 1D**). Similar findings were observed in an *in vivo* model of pancreatic xenograft tumors using CD28-based CAR-T cells against Mesothelin (**Fig S2**). Altogether, these results indicate that chronic antigen stimulation leads to an irreversible T-cell intrinsic program of dysfunction in CD28-based CAR-T cells that causes tumor escape, a process that can be faithfully recapitulated in our *in vivo* models for further mechanistic study.

We next aimed to investigate the molecular basis of CAR-T cell dysfunction. To this end, we performed bulk RNA-sequencing on CD8+ CAR-T cells isolated from SKOV3 tumors during the effective and dysfunctional phase, as well as from the preinfusion product (day 0). Although CAR transcript levels were comparable across groups (**Fig. S3A**), dysfunctional CAR-T cells exhibited widespread transcriptional changes, relative to both effective and preinfusion cells, as revealed by differential expression analysis. Principal component analysis (PCA) demonstrated clear separation between the three conditions, with tight intra-group clustering and no evidence of outliers or marked sample heterogeneity (**Fig. 1F**).

To gain insight into the biological pathways underlying CAR-T cell dysfunction, we performed Ingenuity Pathway Analysis (IPA) on the differentially expressed genes. Dysfunctional CAR-TILs were enriched for gene signatures associated with T cell exhaustion, senescence, and inhibitory checkpoint activation. Natural killer-related genes were also found as top-enriched pathways in dysfunctional CAR-TILs, consistent with previous reports linking NK cell-like transcriptional programs to CAR-T cell dysfunction^14,17^(**Fig.1G and S3B-C**). To benchmark our model against established signatures of T cell dysfunction, we applied the 17-gene exhaustion signature described by Good et al, derived from dysfunctional CD8+ human TILs across four cancer types^14^. All genes in the signature were significantly upregulated in dysfunctional CAR-TILs compared to preinfusion product, suggesting our xenograft model closely mirrored transcriptional features of patient T cell dysfunction. Strikingly, most of these genes were early induced at the effective phase compared to preinfusion (**Fig 1H**), suggesting an early commitment to exhaustion in CAR-TILs. In line with this, broader exhaustion-associated genes sets were also enriched in CAR-T cells shortly after infusion (**Fig. S3D**). Together, these findings indicate that the exhaustion program is initiated early in CAR-T cells following tumor encounter and becomes progressively reinforced under chronic antigen stimulation.

### An *in vivo*-based custom CRISPR screen identifies mediators of CAR-T cell dysfunction

Transcriptomic profiling revealed marked differences between dysfunction and effective CAR-TILs compared to the preinfusion product. Based on these data, we next hypothesized that genes upregulated during the dysfunctional phase contribute functionally to CAR-T cell exhaustion *in vivo*, and that their targeted disruption could delay or prevent dysfunction and thereby enhance antitumor efficacy.

To evaluate these hypotheses, in a first approach we clustered variable genes based on their expression dynamics across the preinfusion, effective and dysfunctional phases of CAR-TILs. This approach identified distinct gene clusters with shared transcriptional temporal trajectories, including groups progressively upregulated during dysfunction and others downregulated from the preinfusion stage. In this analysis, genes associated with T cell exhaustion exhibited upregulation (JUNB, PDCD1, LAG3 and CTLA4, among others), whereas genes linked to T cell memory and proliferation showed progressive downregulation from preinfusion to dysfunctional phase in CAR-TILs (EOMES, CD28 and LIF, among others) (**Fig. S4A**).

We focused on clusters 1, 6 and 7, which contained 1257 genes upregulated during the dysfunctional phase. To prioritize candidates, we first selected genes that were significantly upregulated in the dysfunctional phase compared to the effective phase. These genes were then intersected with previously reported signatures of dysfunctional human TILs and CAR-T cells in cancer^13,14,19–24^. From this intersection, we further prioritized genes based on their inclusion in in vitro CAR-T cell dysfunction signatures ^13,14^ (CST7, MYO7A) or their established roles in T cell regulation (DUSP2, ZFP36), resulting in four candidate genes for further analysis (**Fig. S4B**). We genetically engineered HER2-28z CAR-T cells with individual knockouts of each candidate gene or in the safe harbor site AAVS1 as control, using CRISPR/Cas9 technology. Genetic ablation of candidate genes in CAR-T cells did not improve the antitumor efficacy against ovarian xenograft tumors compared to control (**Fig. S4C-F**).

In a second approach, we conducted a custom *in vivo* CRISPR-based screen. To design the a sgRNA library, we first identified genes upregulated in dysfunctional CAR-T cells by overlapping the differentially expressed gene sets from the three pairwise comparisons (preinfusion vs effective, effective vs dysfunctional, and preinfusion vs dysfunctional) shown in Figure 1F. Based on their temporal expression patterns, we stratified these genes into three groups: (i) early-upregulated genes, induced during the effective phase (early dysfunction), (ii) progressively upregulated genes, showing a continuous increase from preinfusion to dysfunction (progressive dysfunction), and (iii) late-upregulated genes, selectively induced at the dysfunctional stage (late dysfunction) (**Fig. 2A**). We next designed a library of sgRNAs to perform CRISPR/Cas9-mediated knockouts of 300 candidate genes, including the top 100 upregulated genes from early dysfunction, progressive dysfunction and late dysfunction (**Fig 2A**). To screen for modifications that provide an advantage to CAR-T cells *in vivo*, a dose of pooled-KO CAR-T cells was infused to NSG mice bearing SKOV3 tumors. After 3 to 4 weeks of treatment, CAR-TILs were isolated from tumors, and sgRNA abundance was compared between pre-KO T cells and those recovered after the *in vivo* screen (**Fig. S5A**). Per the MAGeCK algorithm, control sgRNA (i.e. non-targeting, intron-targeting and safe harbor-targeting) maintained consistent abundance in CAR-T cells before and after Cas9-induced genetic modifications (**Fig. S5B-D**). In contrast, sgRNAs targeting essential genes for T cell survival, including BCL2, HSPA9, HSPA5, PAFAH1B and CDC16 were significantly depleted during the *in vitro* expansion of genetically modified CAR-T cells (**Fig. S5E and F**). Using this CRISPR screen, we identified 14 significantly enriched genes after chronic antigen stimulation *in vivo*, including ZC3H12C, UCP3, AHNAK2, STON1, MYH6, LILRB1, PTK7, TG, INSRR, ADAM12, COL4A1, ITGB8, FN1 and ACTG2. Notably, these genes showed superior enrichment compared to PD-1 or LAG-3. To validate the *in vivo* screen findings, we selected one top-ranked (ZC3H12C, from early dysfunction), one mid-ranked (TG, from early dysfunction), and one low-ranked significantly enriched gene (ITGB8 from progressive dysfunction), according to the MAGeCK software (**Fig. 2B and Fig S5G**). We next evaluated the impact of ablating ZC3H12C, TG or ITGB8 on the antitumor activity of CAR-T cells using our *in vivo* model of SKOV3 xenograft tumors. CAR-T cells knockedout at AAVS1 safe harbor site were included as controls, and a non-enriched gene KO (RGS2) was added as an additional negative control to test the specificity of the screen hits (**Fig. 2B**).

**Figure 2.**
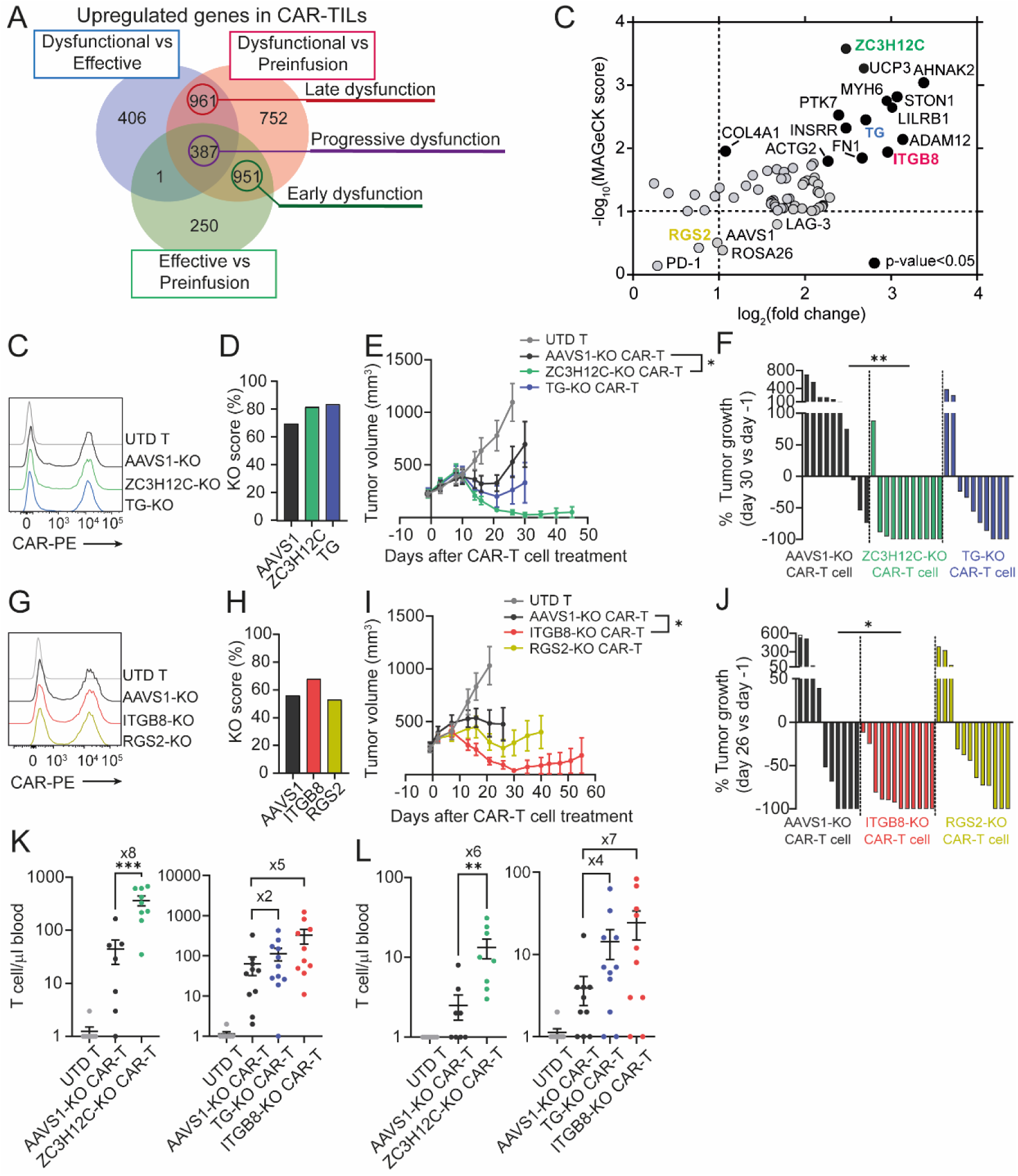
An *in vivo* custom CRISPR-based screen identifies mediators of CAR-T cell dysfunction. **A**) Overlap of upregulated genes across comparisons between preinfusion, effective and dysfunctional CAR-TILs. A custom CRISPR/Cas9 library was generated by selecting 100 genes from early, late, and progressive dysfunction timepoints (indicated by circles). **B**) Gene knockout enrichment following the *in vivo* screen was analyzed from Amplicon-seq data using the MAGeCK algorithm (n=3 healthy donors). The correlation between MAGeCK score and fold change is shown. Significantly enriched genes (MAGeCK score>1 and p-value<0.05) are indicated in black. Candidate genes used for further validation are highlighted. Safe harbour AAVS1 and ROSA26, and positive controls PD-1 and LAG-3 are also depicted. For simplicity of the plot, only genes with a MAGeCK score>1 are shown. **C-F**) NSG bearing pre-established SKOV3 tumors were treated with a single dose of 2×10^6^ untransduced T cells (UTD), anti-HER2-28z CAR+ T cells knocked out for candidate genes ZC3H12C or TG, or AAVS1. **C**) Histogram of CAR expression in CD8+ T cells used for *in vivo* assessment of antitumor effect. **D**) CRISPR/Cas9 efficiency represented as KO score at the end of primary T cell expansion. **E**) Tumor volume of mice treated with KO CAR-T cells was analyzed at indicated timepoints. **F**) Change in tumor volume on day 30 after CAR-T therapy versus baseline is plotted for individual tumors. **G-J**) In a second experiment, the anti-tumor efficacy of ITGB8- or RGS2-KO CAR-T was evaluated as in (C-F). Histogram of CAR expression in CD8+ CAR-T cells used for this experiment (**G**), KO score (**H**), tumor volume of treated mice (**I**) and change in tumor volume at day 26 after CAR-T therapy (**J**). (**E and I**) Data are plotted as mean ±SEM of tumor volume (n=10-12 tumors per group). *p<0.05 by two-way ANOVA with Tukey’s multiple comparisons test at day 26 after CAR-T therapy. (**F and J**) *p<0.05, **p<0.01 by one-way ANOVA at indicated timepoint. **K-L**) Analysis of total T cell concentration in the blood of animals treated with ZC3H12C-KO, TG-KO, ITGB8-KO, control AAVS1-KO CAR-T or UTD T cells (two independent experiments) at day 14 (**K**) and 21 (**L**) after T cell injection. Data are plotted as mean ±SEM. Each dot represents a mouse. Fold change relative to AAVS1-KO group is indicated. **p<0.01, ***p<0.001 by one-way ANOVA with Tukey’s multiple comparisons test.

In a first experiment, HER2-28z CAR-T cells deficient in ZC3H12C or TG were genetically engineered using CRISPR/Cas9 technology. On day 9 after primary T cell activation, all groups showed comparable efficiency of CAR expression and gene editing (**Fig. 2C and D**). We observed that AAVS1-KO CAR-T cells induced only partial responses in mice and in most cases CAR-T therapy failed to control tumor growth. In contrast, ZC3H12C-KO CAR-T cells demonstrated significantly enhanced antitumor effect. Remarkably, eleven out of twelve tumors were completely eliminated after 45 days of treatment with ZC3H12C-KO CAR-T cells. TG-KO CAR-T cells showed a trend toward enhanced antitumor efficacy compared with AAVS1-KO CAR-T cells, although it did not reach statistical significance. TG-KO CAR-T therapy induced a 50% of complete responses in mice with SKOV3 tumors, whereas control AAVS1-KO CAR-T cells only achieved a 10% of complete responses (**Fig. 2E and F**).

In a second experiment, we generated HER2-28z CAR-T cells lacking ITGB8, or RGS2, or AAVS1 as a control (**Fig. 2G and H**). Despite being low ranked in the *in vivo* enrichment screen, knocking-out ITGB8 significantly enhanced the CAR-T cell antitumor efficacy *in vivo* compared to control. Ten out of twelve tumors were completely eliminated in mice treated with ITGB8-KO CAR-T cells (83% of complete responses). In contrast, AAVS1-KO CAR-T induced 50% of complete responses in mice with SKOV3 tumors. The genetic ablation of RGS2 did not enhance the antitumor effect of control CAR-T cells, inducing 42% of complete responses. This was expected since RGS2 was not positively enriched after the *in vivo* screen (**Fig. 2I and J**).

To examine potential correlates of improved antitumor efficacy, we analyzed T cell expansion and persistence in mice with SKOV3 tumors. On day 14 after therapy, CAR-T cells with knockout of target genes (ZC3H12C, TG or ITGB8) showed greater expansion in the blood compared to control, with ZC3H12C-KO CAR-T cells reaching statistically significant higher accumulation (**Fig. 2K**). Although T cell persistence declined in all treated mice by day 21, ZC3H12C-KO CAR-T cells maintained significantly higher T cell numbers, while TG-KO and ITGB8-KO CAR-T cells also trended higher than control AAVS1-KO CAR-T cells (**Fig. 2L**). To test whether these results extend beyond ovarian tumors, we evaluated ZC3H12C in an *in vivo* model of pancreatic xenografts in which CAR-T cells dysfunction also causes tumor escape. Using CD28-based Mesothelin-specific CAR-T cells, disruption of ZC3H12C enhanced both antitumor efficacy and persistence *in vivo,* indicating that its effects extend across tumor types and antigen targets (**Fig. S6**). Altogether, these results highlight our custom *in vivo* functional screen, guided by transcriptomic profiling of CAR-T cell across preinfusion, effective and dysfunctional phases, as an effective strategy for identifying genes that directly contribute to CAR-T cell dysfunction.

### ZC3H12C, TG and ITGB8 are mediators of early dysfunction in CAR-T cells

To determine which CAR-TIL populations express our candidate genes *in vivo*, we performed single-cell RNA and ATAC sequencing on CAR-TILs isolated at effective and dysfunctional phase, and on the preinfusion product. We identified 14 major clusters by Louvain clustering within the samples analyzed (**Fig. 3A**). Clusters 1, 7 and 11 were specific for preinfusion CAR-T cells, while effective CAR-T cells were mostly confined to clusters 3, 5, 9 and 12. Finally, clusters 2, 4 and 14 were associated with dysfunctional CAR-TILs (**Fig. 3B**). ZC3H12C was mostly expressed in effective and dysfunctional, but not in preinfusion clusters (**Fig. 3C**). Using ATAC-seq we studied regulatory changes associated with each cluster. We observed open chromatin sites for ZC3H12C in dysfunctional (2, 4 and 14) but also in effective (3, 5 and 12) clusters. Similar expression patterns were observed for TG and ITGB8 candidates, although their overall expression levels in CAR-TILs were lower (**Fig. S7A and B**). Within the clusters where ZC3H12C was expressed, we detected transcriptionally active chromatin at the PD-1 and TOX loci (**Fig. 3D**). Further, motif analysis of ZC3H12C-positive cells revealed enrichment of transcription factor binding sites previously implicated in T cell exhaustion, including ATF1, JUNB, FOSL2, JUND, and NR4A1 (**Fig. 3E**), thus suggesting a link between ZC3H12C expression and CAR-TILs undergoing exhaustion.

**Figure 3.**
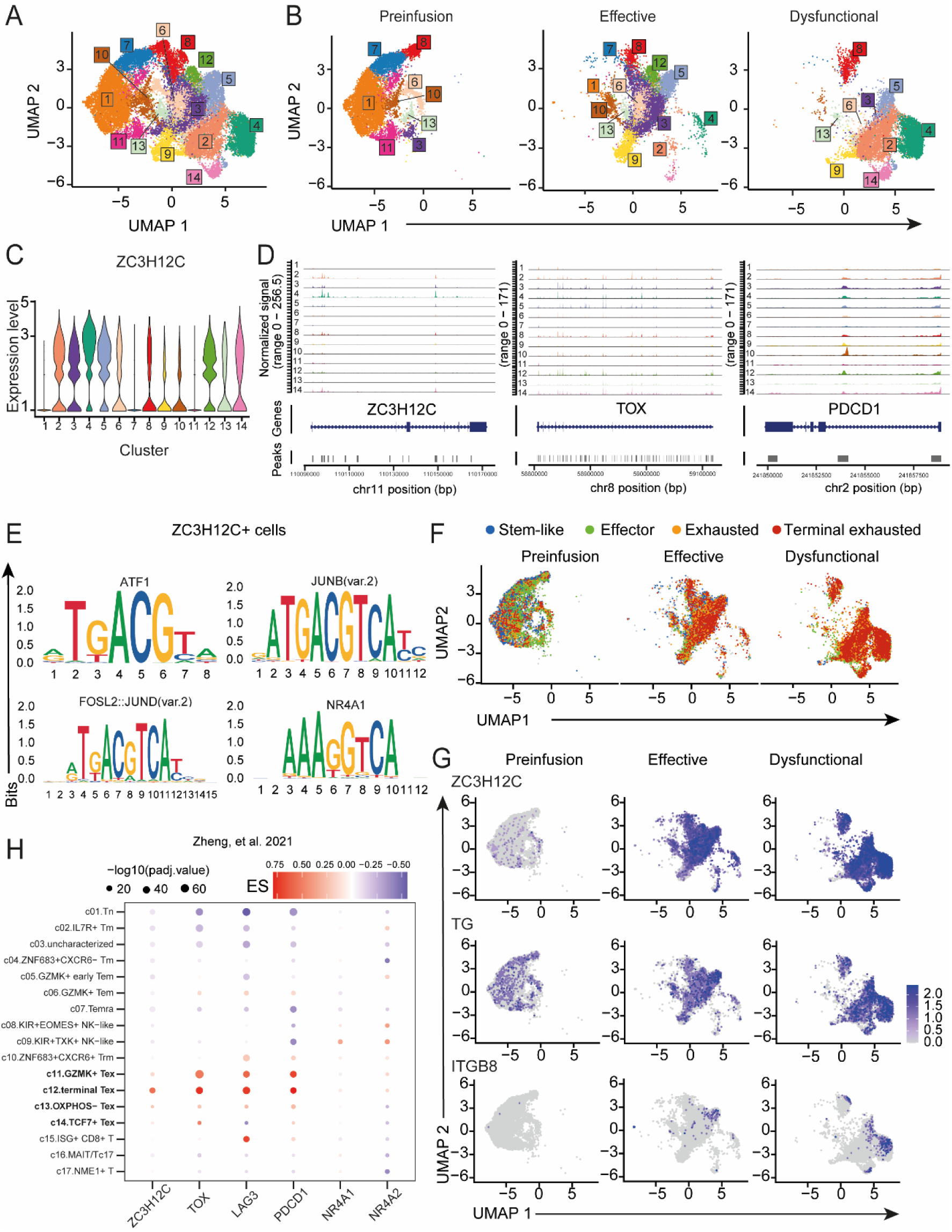
ZC3H12C, TG and ITGB8 are expressed at early phase of exhaustion in CAR-T cells within ovarian xenograft tumors. **A-B**) Uniform manifold approximation and projection (UMAP) plot showing preinfusion CAR-T cells, and effective and dysfunctional CAR-TILs isolated from SKOV3 tumors. Data from one healthy donor independently analyzed. Similar results were observed in three healthy donors. Data are shown with merged timepoints (A) and separated timepoints (B). **C**) Violin plots showing expression of ZC3H12C within clusters identified by single-cell RNA-seq analysis. **D**) Genome browser tracks showing ATAC-seq signal at ZC3H12C, TOX and PD-1 regulatory regions within each of the clusters represented in A. **E**) Motif analysis was performed to identify regulatory elements around the transcription start site (TSS) of differentially expressed genes in ZC3H12C-positive cells. Selected motifs for transcription factors associated with exhaustion from Top24 enriched motifs are shown. **F**) T cell subtype classification integrated with the signature dataset reported by Zheng and colleagues^25^. **G**) UMAP plots of preinfusion CAR-T cells and effective and dysfunctional CAR-TILs color-coded by the density of expression of ZC3H12C, TG or ITGB8. **H**) Dot plot showing expression of ZC3H12C and selected exhaustion-associated genes in meta-clusters identified in CD8+ T cells from Zheng, et al. 2021^25^. Exhaustion clusters are highlighted in bold. Color indicates effect size (ES) and size indicates the level of statistical significance.

To determine the transcriptional phenotype of the CAR-T cell clusters identified in the single cell RNA-seq analysis, we used the signature genes associated to exhaustion, terminal exhaustion, effector and stem-like reported by Zheng and colleagues^25^. This analysis showed that preinfusion CAR-T cells mostly express stem-like and effector markers. These signatures were also identified in effective CAR-TILs. However, at effective phase, we also observed early expression of the exhausted signatures, consistent with our previous findings using RNA-seq analysis (**see Fig. 1H and S1D**). At the dysfunctional phase, CAR-TILs were predominantly identified as exhausted and terminal exhausted T cells (**Fig. 3F and S7C**). ZC3H12C, TG and ITGB8 were most predominantly identified in cells positive for exhaustion signatures. Notably, the three candidate genes were also expressed in effector-annotated cells, but low expression was detected in stem-like positive cells (**Fig. 3G**).

Finally, we validated the transcriptional profiles of our candidate genes in tumor infiltrating T cells derived from cancer patients using the publicly available pan-cancer TIL atlas from Zheng and colleagues, which contains data from 316 patients across 21 cancer types^25^.

We observed that ZC3H12C was expressed in most of the cancer types included in the atlas and its expression was restricted to tumor-infiltrating T cells, when comparing tumor- and blood-derived T cell samples (**Fig. S7D**). Within the 17 meta-clusters describing the pan-cancer T cell atlas, ZC3H12C was found significantly upregulated in the four clusters containing exhausted TILs, while downregulated in clusters related to naïve, memory and effector memory phenotypes in TILs. Similar pattern and level of expression was identified for TOX, PD-1 and LAG-3 in patient-derived TILs. In contrast, other well-established exhaustion-associated factors, including NR4A1 and NR4A2, were less represented than ZC3H12C in patient-derived exhausted TILs (**Fig. 3H**). Similar to ZC3H12C, TG and ITGB8 genes were significantly upregulated in TILs undergoing exhaustion (**Fig. S7E**).

Collectively, these results suggest that candidates ZC3H12C, TG and ITGB8 are associated with the exhausted phenotype in CAR-T cells from our model and in patient-derived tumor infiltrating tumor cells, thus underscoring their potential as targets to prevent CAR-T cell exhaustion at early stages to enhance efficacy against solid tumors.

### ZC3H12C-deficient CAR-T cells demonstrate sustained effector functions after *in vitro* chronic antigen stimulation

We next focused on the role of ZC3H12C, the top candidate identified in the *in vivo* screen, as a mediator of CAR-T cell dysfunction. To characterize the mechanism through which the genetic ablation of ZC3H12C enhances HER2-28z CAR-T cell antitumor efficacy, we performed a repeated stimulation assay *in vitro* where CAR-T cells were chronically exposed to breast cancer tumor cells expressing HER2, to induce T cell dysfunction (**Fig. 4A**). ZC3H12C-KO CAR-T cells showed significantly increased overtime proliferation compared to AAVS1-KO CAR-T cells (**Fig. 4B and S8A**). We also evaluated the cytotoxic capacity of restimulated CAR-T cells using a real-time assay. No differences were detected in cytotoxicity between ZC3H12C-KO and control AAVS1-KO CAR-T cells after acute exposure to target antigen (Day 0). However, while AAVS1-KO CAR-T cells progressively lost their ability to eliminate tumor cells *in vitro* after repeated stimulations, ZC3H12C-KO CAR-T showed significantly improved killing abilities (**Fig. 4C and S8B**). Further, ZC3H12C-KO CAR-T cells showed superior TNF-α, IL-2 and IFNγ production compared to control after several stimulation with tumor cells (**Fig. 4D**). In line with this, when analyzed by intracellular staining, ZC3H12C-KO CAR-T cells showed an increased percentage of polyfunctional T cells producing both IFNγ and TNF-α (**Fig. 4E**). The loss of effector function observed in AAVS1-KO compared to ZC3H12C-KO CAR-T cells during *in vitro* restimulation were T-cell intrinsic, as similar results were obtained when CAR-T cells were stimulated in a CAR-independent manner using PMA/ionomycin following a week of chronic antigen exposure (**Fig. S8C**).

**Figure 4.**
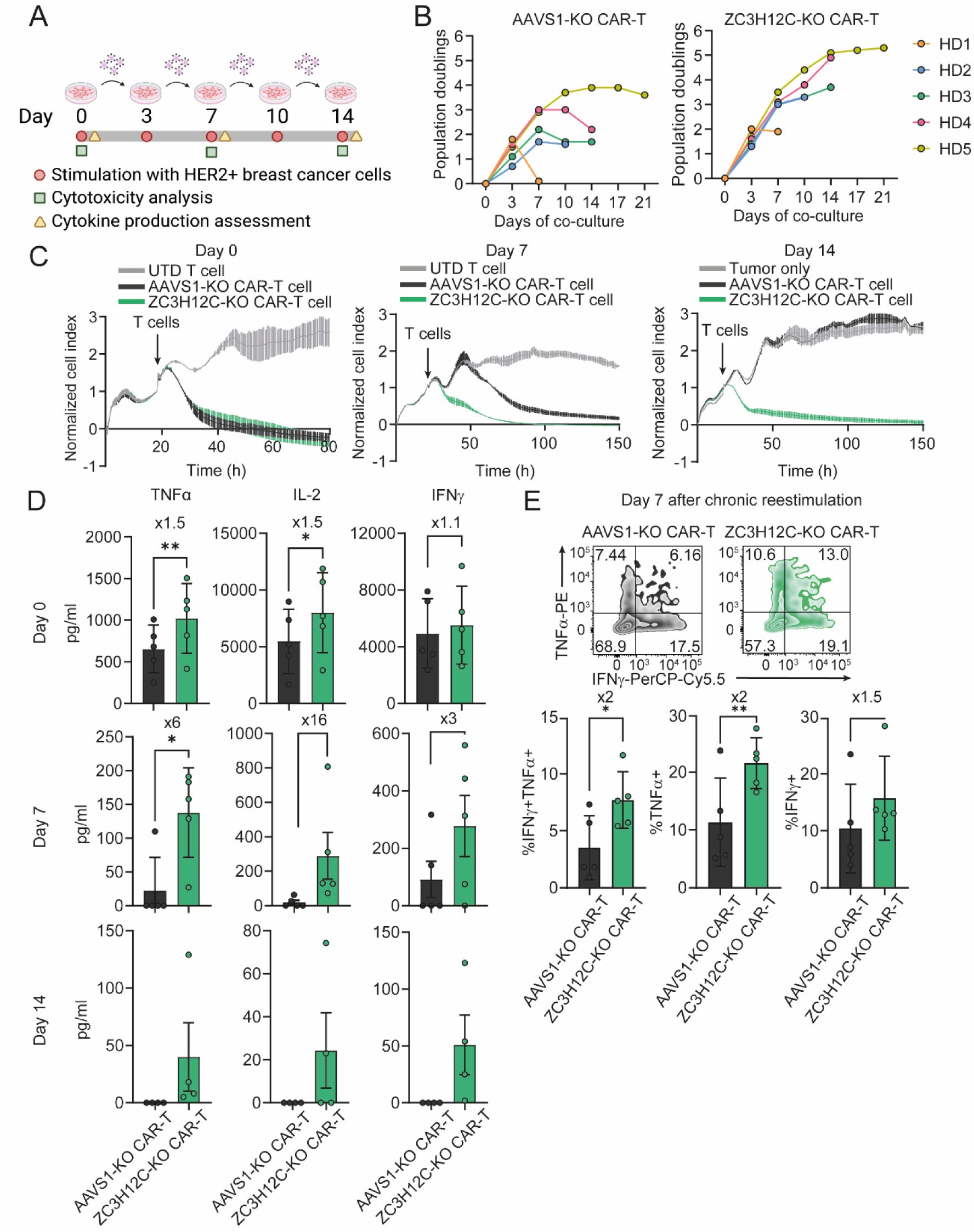
Genetic ablation of ZC3H12C mediates superior effector functions in HER2-28z CAR-T cells after chronic antigen stimulation *in vitro*. **A)** Schematic diagram illustrating the *in vitro* assay used to induce dysfunction through repetitive stimulation of CAR-T cells with antigen. **B**) Population doublings of AAVS1-KO (left panel) and ZC3H12C-KO (right panel) CAR-T cells during *in vitro* restimulation with target cells. Each color represents a different healthy donor. **C**) Cytotoxicity of AAVS1-KO and ZC3H12-KO CAR-T cells against fresh HCC1954 tumor cells (E:T=1:1) was measured at indicated timepoints during the restimulation assay by using a real-time assay. Mean ±SEM of normalized cell index of one representative donor for each timepoint is shown. Time when T cells were added is indicated. **D**) TNF-α, IL-2, and IFNγ production by AAVS1-KO and ZC3H12C-KO CAR-T cells after 24h co-culture with fresh HCC1954 tumor cells (E:T=1:3) was measured at indicated timepoints of *in vitro* restimulation with tumor cells. Data are plotted as mean ±SEM. Each dot represents a healthy donor (n=5 for day 0 and day 7; n=4 for day 14). Fold increase of cytokine secretion by ZC3H12C-KO versus AAVS1-KO CAR-T cells is indicated. *p<0.05, **p<0.01 by two-tailed paired T-test. **E**) Representative flow cytometry plots of intracellular TNF-α and IFNγ staining in AAVS1-KO and ZC3H12C-KO CAR-T cells (gated on live/CD45+) after 24h co-culture with fresh SKOV3 cells (E:T=1:3) on day 7 after repetitive antigen stimulation *in vitro* (top). Frequencies of single and double-positive T cells for IFN-γ and TNF-α are represented as absolute numbers (bottom). Data are plotted as mean ±SD. Each dot represents a healthy donor (n=5). *p<0.05, **p<0.01 by two-tailed paired T-test. Fold change of ZC3H12C-KO relative to AAVS1-KO is indicated (bottom).

### The genetic ablation of ZC3H12C prevents CAR-T cell exhaustion in an *in vivo* model of solid tumors

To assess whether the genetic ablation of ZC3H12C prevents CAR-T cell dysfunction *in vivo*, mice bearing SKOV3 tumors were treated with AAVS1- or ZC3H12C-KO CD28-based HER2-CAR-T cells. After two or three weeks of treatment, CAR-TILs were isolated to analyze phenotype and *ex vivo* function (**Fig. 5A**). We observed superior T cell infiltration in tumors treated with ZC3H12C-KO CAR-T cells compared to control, suggesting that knocking-out ZC3H12C enhances CAR-T cell proliferation *in vivo* (**Fig. 5B**). ZC3H12C-KO CAR-TILs showed reduced expression of exhaustion markers PD-1, TIM-3 and LAG-3, compared to AAVS1-KO CAR-TILs (**Fig. 5C**). ZC3H12C-KO CAR-TILs maintained the ability to kill SKOV3 tumor cells *ex vivo*. In contrast, AAVS1-KO CAR-TILs showed impaired cytotoxic capacity (**Fig. 5D**). In line with this, ZC3H12C-KO CAR-TILs produced significantly higher levels of effector cytokines compared to control, including TNF-α, granzyme B, perforin, IFNγ and IL-6 (**Fig. 5E**). Similar differences were observed after CAR-independent stimulation of CAR-TILs (**Fig. S8D**).

**Figure 5.**
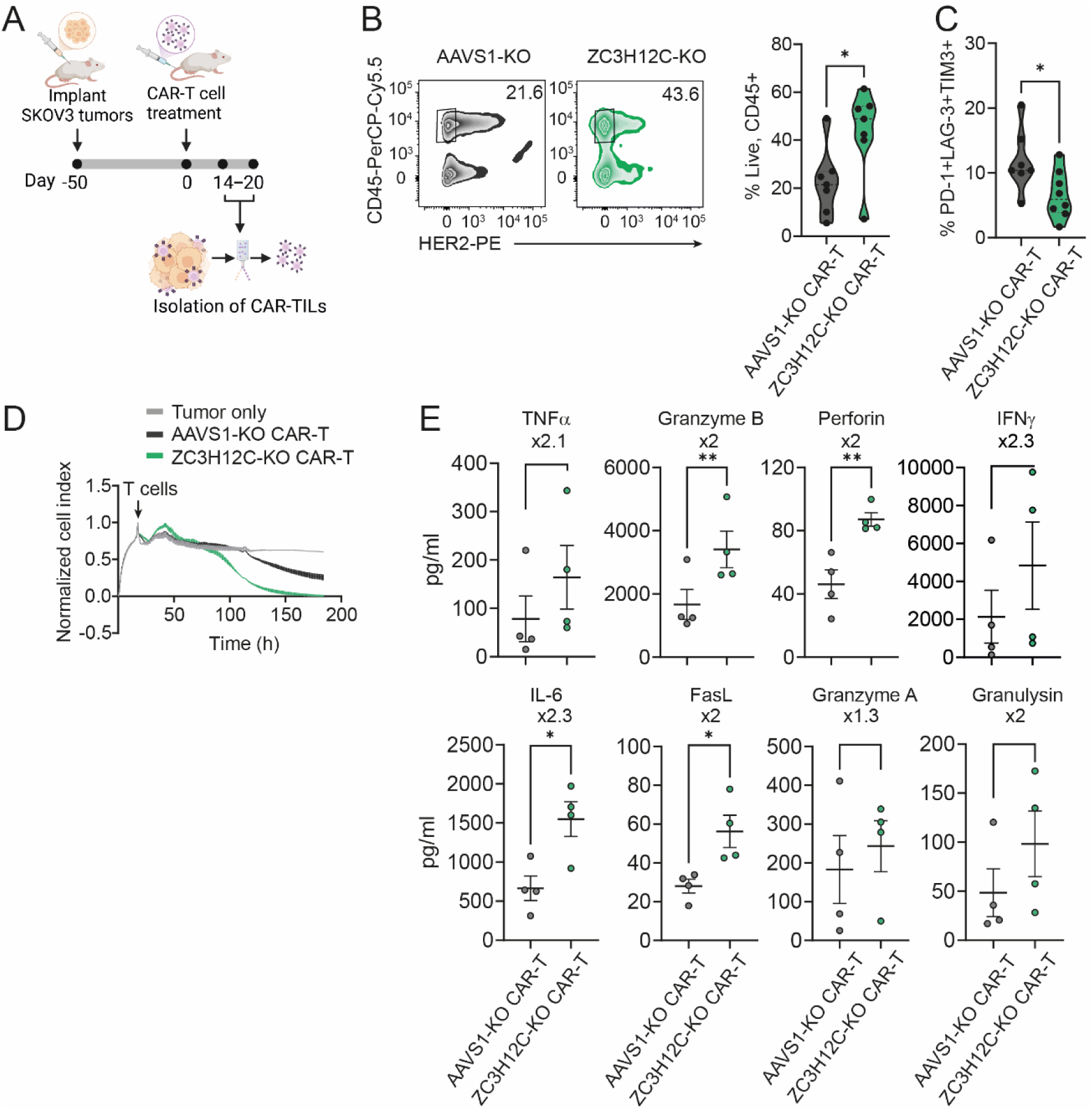
The ablation of ZC3H12C prevents HER2-28z CAR-T cell exhaustion in an *in vivo* model of ovarian xenograft tumors. **A**) Schematic representing the experimental approach used to analyze the *ex vivo* function of ZC3H12C-KO CAR-TILs. **B**) Analysis of CD8+ T cell infiltration in tumors treated with AAVS1-KO or ZC3H12C-KO CAR-T cells on day 17 after infusion. Representative flow cytometry plots showing CD45 and HER2 staining after excluding dead cells (left panel). Violin plots showing the frequency of live/CD45+ cells in tumors treated with AAVS1-KO or ZC3H12C-KO CAR-T cells (right panel). Each dot represents a tumor (n=2 healthy donors, 7-8 tumors per group). Fold change relative to AAVS1 is indicated. **C**) Frequency of PD-1+ LAG-3+ TIM3+ CD8+ CAR-TILs (gated as live, CD45+). *p<0.05 by two-tailed unpaired t-test. **D**) *Ex vivo* cytotoxicity of AAVS1-KO and ZC3H12C-KO CAR-TILs was analyzed against SKOV3 tumor cells (E:T=5:1) using a real-time assay. Time when T cells were added is indicated. Data are plotted as mean ±SEM of normalized index of two duplicates (n=1). **E**) Cytokine production by AAVS1-KO and ZC3H12C-KO CAR-TILs after a 24h *ex vivo* co-culture with SKOV3 tumor cells (E:T=3:1). Data are plotted as mean ±SEM. Each dot represents a healthy donor (n=4). *p<0.05, **p<0.01 by two-tailed paired t-test. Fold change of ZC3H12C-KO relative to AAVS1-KO is indicated.

### Disruption of ZC3H12C enhances antitumor effect and persistence of 4-1BB-based CAR-T cells

We next aimed to determine whether the improved antitumor efficacy observed when knocking-out ZC3H12C in CD28-based CAR-T cells could also apply to CAR constructs with a different co-stimulatory signaling domain. To test this, we used ARI-0001 CAR-T cells, an anti-CD19 academic CAR-T product developed by Hospital Clínic-IDIBAPS, which contains 4-1BB costimulatory domain^26^, to target a newly developed artificial model of aggressive lung xenograft tumors overexpressing CD19.

While all animals treated with AAVS1-KO ARI-0001 showed progressive disease, treatment with ZC3H12C-KO CAR-T cells induced complete responses in all treated animals, with all responses maintained overtime (**Fig. 6A-B**). In line with this, ZC3H12C-KO CAR-T cells showed significantly greater expansion and persistence compared to AAVS1-KO CAR-T cells on days 14 and 21 after CAR-T cell infusion (**Fig. 6C**).

**Figure 6.**
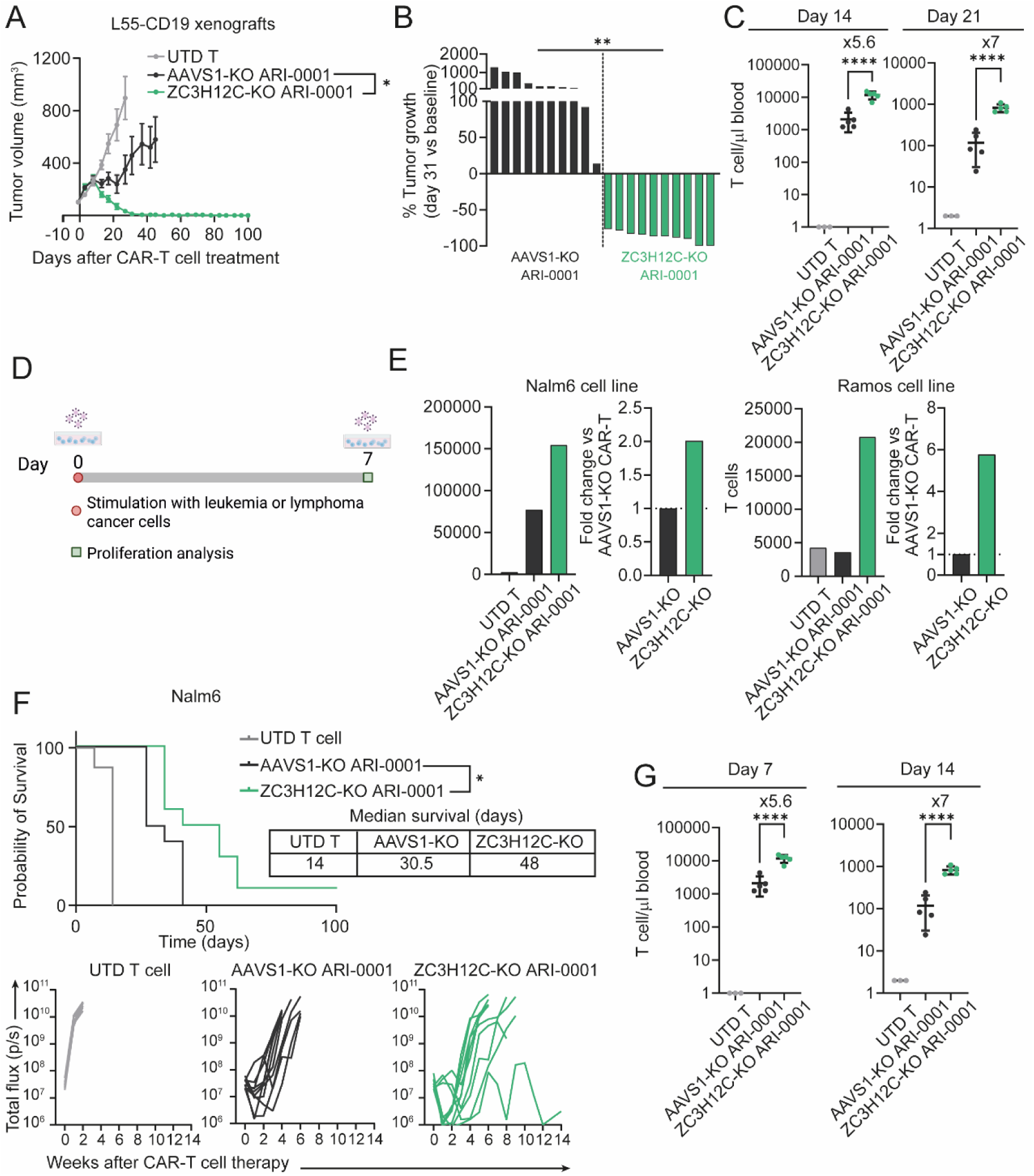
Genetic ablation of ZC3H12C improves CAR-T antitumor efficacy against solid and hematological malignancies using a 4-1BB-based CAR (ARI-0001). **A-C**) NSG mice bearing subcutaneous L55-CD19 tumors were treated with a single dose of 4×10^6^ control unstransduced (UTD) T cells, AAVS1-KO or ZC3H12C-KO CAR+ T cells. **A**) Tumors were measured at indicated timepoints. Data are plotted as mean ±SEM of tumor volume (n=8-10 tumors per group). *p<0.05, by two-way ANOVA with Tukey’s multiple comparisons test at day 27. **B**) Change in tumor volume on day 31 after CAR-T cell therapy. **p<0.01 by two-tailed unpaired t-test. **C**) The concentration of total T cells was determined in the blood of animals treated with ZC3H12C-KO, AAVS1-KO ARI-0001 CAR-T cells, or UTD T cells, at day 14 (left panel) and day 21 (right panel) after treatment. Data are plotted as mean ±SD. Each dot represents a mouse (n=1 healthy donor, 8-10 tumors per group). ****p<0.0001 by one-way ANOVA with Tukey’s multiple comparisons test. Fold change relative to ZC3H12C-KO group is indicated. **D**) Schematic diagram illustrating the in vitro assay using hematological cancer cells. **E**) T cell proliferation after an *in vitro* 7-day co-culture of CAR-T cells with NALM-6 or Ramos cell lines (E:T=1:2) analyzed by flow cytometry. For each cell line, T cell is represented as absolute numbers (left panel) or fold change compared to AAVS1-KO CAR-T cell group (right panel) (n=1 healthy donor). **F-G**) NSG mice were intravenously injected with Nalm6-GFP-luciferase cells. After 7 days, mice were randomized and treated with 2×10^6^ CAR+ T cells, including UTD control T cells, AAVS1-KO or ZC3H12C-KO ARI-0001 (n=10 mice per group, 2 healthy donors). **F**) The Kaplan-Meier survival curves of treated mice are plotted and median survival days calculated. *p<0.05 by Log-rank (Mantel-Cox) test. **G**) T cell persistence in the blood of treated animals was determined on days 7 and 14 after CAR-T cell therapy. Each dot represents a mouse. *p<0.05 by one-way ANOVA with Tukey’s multiple comparisons test. Fold change relative to ZC3H12C-KO group is indicated.

Finally, we evaluated the potential of ZC3H12C-KO ARI-0001 to improve the responses against hematological malignancies. *In vitro*, ZC3H12C-KO ARI-0001 showed superior proliferation after 7 days in co-culture with leukemia and lymphoma cell lines compared to control AAVS1-KO (**Fig. 6D and E**). *In vivo*, treatment with ZC3H12C-KO ARI-0001 resulted in significantly prolonged survival compared to the AAVS1-KO ARI-0001 group. While most animals treated with ARI-0001-KO were sacrificed by week 5, treatment with ZC3H12C-KO slowed disease progression, with 50% of the animals surviving up to week 8 (**Fig. 6F**). In line with these results, the KO of ZC3H12C induced significantly enhanced T cell expansion on day 7 after CAR-T cell therapy compared to control. Similar differences were observed on day 14 after treatment (**Fig. 6G**). Together, the data demonstrate that the functional enhancement observed in ZC3H12C-deficient CD28-based CAR-T cells are generalizable to CAR-T cells containing the 4-1BB costimulatory domain and indicate that the disruption of ZC3H1C could enhance the efficacy of CAR-T cell therapy against both solid and hematological malignancies.

## Discussion

CAR-T cell dysfunction following prolonged exposure to antigen within tumors represents a key factor contributing to the limited efficacy of CAR-T therapy in patients with solid malignancies. However, the mechanisms accounting for this have not been studied in the context of *in vivo* models. Here, we performed a custom CRISPR-based screen using a model of xenograft tumors that recapitulates tumor escape due to CAR-T cell intrinsic dysfunction. Using this strategy, we identified 14 potential novel targets to enhance CAR-T cell efficacy against solid malignancies by preventing T cell dysfunction. Genetic ablation of the top candidate ZC3H12C enhanced CAR-T cell functional persistence and mediated superior antitumor responses against xenograft tumors.

*In vitro* models have provided insights into the mechanisms behind the loss of CAR-T cell functions. Some studies have used tonic signaling CARs that exhibit constitutive, antigen-independent activation of downstream signaling pathways, leading to the premature acquisition of an exhausted phenotype^13,15^. In others, CAR-T cells have been repetitively stimulated with tumor cells to induce sustained antigen exposure. The loss of functional properties in this setting is in part dependent on the surface expression of the CAR and can be largely reversed after a resting period in the absence of tumor cells, allowing CAR re-expression on the cell surface^14,16,17^. Consequently, *in vitro* models may not fully capture the mechanisms driving the loss of CAR-T cell functions.

Here, we established an *in vivo* model in which chronic antigen exposure drives progressive loss of CAR-T cell functions that was refractory to restoration, due to T cell intrinsic mechanisms. Transcriptomic characterization revealed that exhaustion signatures characteristic of the transcriptome of CAR-TILs at dysfunctional phase are also found in CAR-TILs at effective phase. This suggests that although infused CAR-T cells elicited significant antitumor responses in mice, they are committed to the exhaustion program early after tumor recognition, ultimately causing therapy failure. Our results are aligned with previous by Schietinger and colleagues in endogenous T cells from a syngeneic model of cancer suggesting that while early T cell dysfunction is a reversible, plastic differentiation state in which effector functions can be restored, it ultimately evolves to a fixed state. T cells committed to late-stage dysfunction could not be rescued by antigen withdrawal or checkpoint blockade^27,28^. Recent studies indicate that in the context of large, pre-established tumors, the exhaustion program is triggered in T cells within hours after initial tumor encounters and activation^29^. From the 14 enriched genes identified in this study, 10 were already upregulated at early stages after treatment, including ZC3H12C, TG and ITGB8. By disrupting these genes, we showed that it is possible to prevent the progression to late-stage dysfunction and induce long-lasting CAR-T cell-mediated antitumor responses.

In this study, we followed two orthogonal discovery routes to identify the mediators of CAR-T cell dysfunction using RNA-seq data obtained from our model. First, we prioritized candidate genes that were significantly upregulated in the dysfunctional phase compared to the effective phase and based on their inclusion in previously published transcriptional signatures of dysfunctional human CAR-T cells and TILs from cancer patients^19–24^. Despite being strongly associated with T cell exhaustion, the genetic ablation of the four candidates tested failed to provide an advantage to CAR-T cell-mediated antitumor effect. Among them, we evaluated ZFP36. Results reported here are aligned with previous findings showing that targeting ZFP36 is insufficient to enhance the antitumor efficacy of CAR-T cells against solid xenografts^30^. By contrast, our in vivo CRISPR-based functional screen identified 14 candidate genes, three of which were individually validated and all enhanced CAR-T cell performance in mice bearing ovarian xenografts. Interestingly, these three genes were upregulated early after T cell activation, a stage that was not captured by our first, transcriptionally guided approach. This discrepancy highlights a key limitation of our first approach: by filtering only for late-stage upregulation, we inadvertently missed critical early drivers of CAR-T cell dysfunction. Together, these findings underscore that CAR-T cell exhaustion is a complex, multifactorial process involving overlapping and potentially redundant mediators, and they emphasize that functional screening approaches are essential for accurately identifying the genes that causally regulate CAR-T cell dysfunction.

Genome-wide loss-of-function libraries have proven successful for the identification of genes regulating immune functions of human T cells^31,32^. Despite their strong potential to uncover unexpected genes mediating a specific phenotype due to their unbiased methodology, genome-wide screens require large number of cells to maintain library representation. This is particularly challenging with primary or patient samples, or for *in vivo* experiments, where only limited numbers of cells can be administered to and recovered from mice with xenograft tumors. To overcome this limitation, Knudsen et al., restricted their library to 135 targets with previously characterized roles in T cell function and conducted an *in vivo* screen in CAR-T cells^33^. Similarly, Datlinger, et al. performed initial genome-wide screens in CAR-T cells *in vitro*, from which they prioritized 39 top candidates for subsequent *in vivo* screening^34^. Although narrower in scope than genome-wide screens, these approaches demonstrate the feasibility and utility of *in vivo* screens for identifying regulators of CAR-T cell function. However, prior studies have focused on models of hematological malignancies. Therefore, the factors contributing to CAR-T cell dysfunction in solid tumors remain largely unexplored. Here, we conducted a CRISPR-based screen targeting 300 of the most highly upregulated genes identified during *in vivo* CAR-T cell dysfunction. Among the 14 enriched candidates, all three that were functionally validated mediated superior CAR-T cell persistence and enhanced antitumor response. These results validate the effectiveness of our strategy and lead us to hypothesize that the remaining candidates may likewise contribute to enhance CAR-T cell efficacy by preventing dysfunction against solid malignancies.

ZC3H12C gene, which encodes for the CCCH zinc finger protein Regnase-3 (also known as MCPIP3), appeared as top hit in our *in vivo* screen. Regnase proteins have been identified as regulators of the immune response by directly modulating the expression of proinflammatory cytokines through their RNA-binding activity and endonuclease domain^35^. While Regnase-3 has been characterized to restrict the expression of TNF-α and IL-6 in myeloid cells^36^, its role in regulating effector functions of human T cells has not been described. In this study, we observed that knocking-out ZC3H12C in CAR-T cells induces increased proliferation and superior production of effector cytokines, leading to effective, fast elimination of xenograft tumors and preventing the acquisition of an exhausted phenotype in CAR-TILs after chronic antigen stimulation. Further research is required to identify the molecular targets of Regnase-3 in T cells. In previous studies, two additional CCCH zinc finger proteins, Regnase-1 (ZC3H12A gene) and Roquin-1, were identified through separated CRISPR screens using murine OT-I cells to enhance T cell expansion and antitumor functions in mouse models^37,38^. More recently, a study using human T cells showed that dual ablation of Regnase-1 and Roquin-1 improved the efficacy of adoptive T cell therapy against solid tumors, albeit with associated toxicities in mice^39^. In our model, ZC3H12A and Roquin-1 were significantly upregulated in CAR-TILs at dysfunctional phase but were not identified within the top upregulated genes and therefore were not included in our CRISPR-based screen. Next studies will be required to elucidate the contribution of the different members of the Regnase family in regulating CAR-T cell function.

A main limitation for this study is the use of an immunodeficient mouse model, which lacks the immunosuppressive networks within the tumor microenvironment (TME) that can also drive T cell dysfunction^40^. However, the similarities observed between the signatures of dysfunctional CAR-TILs described here and those previously reported from dysfunctional TILs in cancer patients^25^ indicate that our candidate genes may be of significance in the context of an intact TME. Importantly, the expression of ZC3H12C, TG and ITGB8 in exhausted T cells was confirmed in single-cell RNA-sequencing data from cancer patients across multiple cancer types. We hypothesize that the improved intrinsic fitness of ZC3H12C-KO CAR-T cells could enhance their persistence and function in the presence of immunosuppressive pathways. In addition, the increased secretion of proinflammatory cytokines by ZC3H12-KO CAR-T cells may help reshape the tumor milieu into a more proinflammatory state, thereby amplifying immune responses against tumor cells. Of note, in a complementary study submitted alongside the present work, Kavishwar *et al.* independently identified ZC3H12C expression as a feature of exhausted T cells through analysis of molecular signatures in patient-derived TILs. Their study revealed specific chromatin remodeling at the ZC3H12C locus, accompanied by increased expression as TILs progressed toward a dysfunctional state. This induction was restricted to T cells undergoing chronic stimulation and was not observed during acute T cell responses. Moreover, they demonstrated that genetic ablation of ZC3H12C similarly enhances the persistence and antitumor efficacy of both CAR- and TCR-engineered T cells *in vivo*. Together, these complementary findings establish ZC3H12C as a conserved regulator of T cell dysfunction across adoptive T cell platforms.

In conclusion, through an *in vivo* CRISPR screen, we identified novel and promising gene targets for engineering CAR-T cells with enhanced resistance to exhaustion. The potent antitumor efficacy of ZC3H12C-deficient CAR-T cells observed across tumor models strongly support the translation of these findings into clinical testing.

## Materials and Methods

### Experimental Animals

All mouse studies were conducted under protocols (184/20 and 238/21) approved by the Ethics Committee for Animal Experimentation (CEEA) of the University of Barcelona and the Generalitat de Catalunya. Mice were bred and housed at the University of Barcelona’s Animal Facility.

### Mouse xenograft study

For solid tumor models, NSG mice, either female (studies with SKOV3 tumors) or male (studies with L55 or CAPAN-2) were subcutaneously implanted with 5×10^6^ tumor cells in one or both flanks, in a 50% solution of Matrigel (Corning) in phosphate-buffered saline (PBS). Prior to treatment, mice were randomized based on tumor volume. CAR-T cells were intravenously administered to tumor-bearing mice in doses of 1×10^6^-1×10^7^ CAR+ T cells per animal in 100 µl of PBS (CAR-T cell dose and tumor volume at treatment are indicated in figure legends). Tumor dimensions were measured every 4-5 days using a digital caliper and tumor volumes were calculated using the formula: V = 6 x (L x W2) / π, where L is length and W is width of the tumor. Mice were sacrificed when tumors reached 1500 mm^3^.

For leukemia models, male NSG mice were intravenously injected with 1×10^6^ Nalm6 tumor cells expressing GFP-FFLuc. One week later, mice were randomized and intravenously treated with 2×10^6^ CAR+ T cells per animal in 100 µl of PBS. Disease progression was weekly monitored using bioluminescence with Xenogen IVIS 200 Imaging System (PerkinElmer). Data analysis was performed using Living Image software (PerkinElmer). Mice were sacrificed when reached a total flux value of 1×10^10^ photons per second (p/s).

T cell persistence was analyzed from peripheral blood samples obtained by retro-orbital bleeding at indicated timepoints. Blood samples were mixed with antibodies against T cell markers CD45, CD4 and CD8 (table 1) in BD Trucount tubes (BD Biosciences, # 340334), and the absolute number of T cells per microliter was calculated following manufacture’s protocol.

**Table 1.** List of antibodies and other reagents used for flow cytometry and cell sorting.

| Antibody | Reference |
| --- | --- |
| Anti-human CD45 (HI30) PerCP-Cy5.5 | BD Biosciences, #564105 |
| Anti-human CD45 (HI30) APC | Biolegend, #304012 |
| Anti-human CD4 (RPA-T4) AF488 | BD Biosciences, #557695 |
| Anti-human CD4 (OKT4) PE | Invitrogen, #12-0048-42 |
| Anti-human CD8 (HIT8a) APC | Biolegend, #300912 |
| Anti-human CD8 (SK1) APC-H7 | BD Biosciences, #566856 |
| Anti-human PD-1 (EH12.1) PeCy7 | BD Biosciences, #561272 |
| Anti-human PD-1 (EH12.2H7) BV711 | Biolegend, #329928 |
| Anti-human LAG-3 (11C3C65) BV605 | Biolegend, #369316 |
| Anti-human LAG-3 (3DS223H) PeCy7 | Invitrogen, #25-2239-42 |
| Anti-human TIM3 (F38-2E2) PerCP-eFluor710 | Invitrogen, #46-3109-42 |
| Anti-human CD39 (A1) PE-Dazzle | Biolegend, #328224 |
| Anti-human Ki67 (Ki-67) BV605 | Biolegend, 350522 |
| Anti-human TNF-α (MAb11) PE | BD Biosciences, #554513 |
| Anti-human IFNγ (B27) PerCP-Cy5.5 | BD Biosciences, #560704 |
| Anti-human HER2 (24D2) PE | Biolegend, #324406 |
| Streptavidin PE | Thermo Fisher #12-4317-87 |
| Streptavidin APC | Invitrogen, #17-4317-82 |
| Streptavidin eFluor450 | Thermo Fisher, #48-4317-82 |
| Fixable viability dye eFluor™ 450 | eBioscience, #65-0863-14 |
| FcR blocking reagent mouse | Miltenyi, #130-092-575 |

### CAR construction and lentiviral production

The single-chain variable fragments (scFv) used for targeting HER2 with low affinity (4D5.5) and human mesothelin (M11) were was previously described^18,41^. The scFv sequence targeting CD19 (FMC63) was retrieved from GenBank identifier HM852952.1. These scFvs were used in a second-generation CAR with the CD8 hinge, the CD28 transmembrane and intracellular domains and the CD3ζ signaling domains. ARI-0001 CAR contains the CD19-binding scFv A3B1, a CD8α extracellular hinge region and transmembrane domain, along with the 4-1BB costimulatory and CD3ζ endodomains^26^.

The sequence of truncated CD19 (CD19t) corresponds to NM_001770 variant 2 (GeneBank), including the extracellular and transmembrane domains, and 4 amino acids of the cytoplasmatic tail of the protein sequence^42,43^. Similarly, the sequence of truncated HER2 (HER2t) consists on the extracellular and transmembrane domains of the aminoacidic sequence P04626.

All constructs, including CARs and truncated proteins, were synthesized by BaseClear B.V. (Leiden, Netherlands) or GenScript Biotech (New Jersey, United States of America) and cloned into a third-generation lentiviral vector pCCL under the control of EF1a promoter^26^.

Lentiviral vectors were generated by transfecting HEK293FT cells and tittered as previously described^44^. Briefly, 10×10^6^ HEK293FT cells were seeded in a total volume of 18 mL of medium in a p150 culture plate. Eighteen hours later, cells were transfected with 18 µg of transfer vector (containing CAR) and a pre-mixed packaging mix containing 15 µg of pREV, 15 μg of pRRE and 7 μg of pVSV, using PEI (Polysciences, #23966-100). The lentivirus-containing supernatant was collected 48 and 72h after transfection, 0.45 µm filtered, and concentrated using Lenti-X Concentrator (TakaraBio, #631232) as per manufacturer’s instructions. Lentivirus were stored at −80°C until use. Lentiviral vectors were tittered based on CAR expression in the T cell membrane using wild-type Jurkat cells thought the limiting dilution method.

### Cell line culture

All human cancer cell lines were purchased from the American Type Culture Collection (ATCC) except for HEK 293 FT (human embryonic kidney cels), which was obtained from ThermoFisher (#R70007) and L55 (non-small cell lung cancer), which was kindly provided by Dr. Carl June at University of Pennsylvania. HEK 293FT were cultured in DMEM (Dubbeco’s Modified Eagle’s Medium high glucose, Gibco, #10741574) supplemented with 1% GlutaMax (ThermoFisher Scientific, #35050-061) and 1% non-essential amino acids (NEAA) (ThermoFisher Scientific, #11140-061). SKOV3 (ovarian cystadenocarcinoma, #HTB-77) were cultured in DMEM. CAPAN-2 (pancreatic carcinoma, #HTB-80) were cultured in DMEM/F12 (Gibco, #11320033). HCC1954 (breast ductal carcinoma, #CRL-2338), L55, Ramos (Burkitt’s lymphoma, #CRL-1596) Nalm6 (acute lymphoblastic leukemia, #CRL-3273) and Jurkat clone E6.1 (acute T-cell leukemia, #88042803-1VL) were cultured in RPMI-1640 (Roswell Park Memoria Institute, Gibco, #11530586). All media were supplemented with 10% FBS (Merck, #9665) and with penicillin-streptomycin (P/S) (100mg/mL) (Invitrogen, #15140122). Cells expressing CD19, HER2 or GFP-FFLuc were generated by transducing the wild type cell lines with lentiviral vectors expressing CD19t, HER2t (truncated version of CD19 or HER2, respectively) or GFP-FFLuc. All cell lines were maintained in a humidifier at 37 °C and 5% CO_2_ and regularly validated to be *Mycoplasma* free.

### Isolation, transduction, electroporation and expansion of human primary T cells

Human T cells were isolated from healthy donor buffy coats obtained from the Barcelona Public Blood and Tissue Bank and CAR-T cells were generated and expanded as previously described^41^. Briefly, CD4 or CD8 lymphocytes were separated using RosetteSep for human T cells (Stem Cell Technologies, #15062 and 15063) and stimulated with CD3/CD28 human T-Activator Dynabeads (Gibco, #11132D). After 24h of activation, T lymphocytes were transduced with CAR-encoding lentivirus at indicated MOI. Activation beads were removed on day 3 after isolation. Then, T cells were daily counted from day 5 to day 10 after isolation and maintained at a concentration of 0.8 ×10^6^ cells/mL. On day 10, T cells were frozen in aliquots containing a 1:1 ratio of CD4 and CD8 T cells and stored within a liquid nitrogen tank.

T lymphocytes were cultured in complete RPMI 1640 media containing 10% heat-inactivated fetal bovine serum (FBS) (Merck, #F4135), 1% Penicillin-Streptomycin (ThermoFisher, #15140122), 1% HEPES (Sigma-Aldrich, #H3375-25G), 1% GlutaMax (ThermoFisher, #35050-061) and 100 or 5 IU of IL-2 (Proleukin^®^) for CD8 and CD4 T cells, respectively. CRISPR-modified CAR-T cells were expanded in the presence of 10ng/mL human IL-7 (Miltenyi Biotec, # 130-095-362) and IL-15 (Miltenyi Biotec, # 130–095-764) to enhance gene editing.

To generate a pool of KO CAR-T cells, T cells were cotransduced with two lentiviral vectors: i) a lentiviral vector encoding the HER2-28z CAR and ii) a pool of lentiviral vectors at a MOI of 0.2-0.5 to ensure that T cells were transduced with only one vector containing a sgRNA. On day 4 after isolation, T cells were debeaded and GFP-positive cells (as a surrogate of LV integration for the library) were separated by FACS-sorting. After sorting, T cells were rested for 2 h and subsequently electroporated with Cas9 protein. The T-cell expansion protocol was then continued as aforementioned from day 5 to day 10-11.

### CRISPR/Cas9-mediated gene editing in human T cells

To generate a pool of KO CAR-T cells using a custom library of sgRNAs, we electroporated Cas9 protein to library-expressing CD4+ and CD8+ CAR-T cells at day 4 after activation using Neon transfection system (ThermoFisher). Briefly, 4-7×10^6^ library-expressing CAR-T cells were washed with PBS and resuspended in 60 µl of Buffer R and kept on ice. 10 μg of TrueCut^TM^ Cas9 Protein v2 (ThermoFisher Scientific, #A36499) protein were then mixed with 60 μL of Buffer R and added to T cells. 100 μL of the mixture were pipetted with the Neon™ Transfection System Pipette (ThermoFisher Scientific, #MPP100) and electroporation protocol #24 (1600V, 10ms, 3 pulses) was performed in the Neon™ Transfection System Pipette Station (ThermoFisher Scientific, #MPS100). After electroporation, cells were immediately transferred to tissue culture plates containing conditioned medium (T cell culture media recovered from the supernatant after the first centrifugation) at 1.5×10^6^ cells/mL for expansion.

To knockout ZC3H12C, TG, ITGB8, RGS2 or AAVS1 in both human CD8+ and CD4+ human T cells we used Lonza™ P3 Primary Cell 4D-Nucleofector™ X Kit L (Lonza™ V4XP-3024) with TrueCut^TM^ Cas9 Protein v2 (ThermoFisher Scientific, #A36499) protein and a chemically synthesized sgRNA targeting each candidate gene (AAVS1: GGGGCCACUAGGGACAGGAU^13^, ZC3H12C: UGUGUCCAAUGAUAACUACA, TG: GGCCUGGUCACAUUGCACUG, ITGB8: AGCACAUGGAUGUAUCCAUG, or RGS2: ACTCCTGGGAAGCCCAAAAC). sgRNAs targeting candidate genes were selected from the custom CRISPR-based library and purchased from Synthego. In brief, ribonucleoprotein complex was mixed at sgRNA:Cas9 molar ratio of 3:3:1 in a final volume of 10 µL of P3 buffer and incubated for 20 min at RT. 4-6×10^6^ T cells were washed with PBS and resuspended in 90 µL of P3 buffer. Cell suspensions were then combined with newly formed RNP complexes and immediately transferred to a Nucleocuvette. Electroporation protocol EH-115 was performed on a Nucleofector Unit. After electroporation, 100 µL of warm media was added to the Nucleocuvette and cells were allowed to recover for 15 min in the incubator, before being transferred into tissue culture plates at 1.5×10^6^ cells/mL with conditioned medium for expansion. To assess the efficiency of CRISPR/Cas9-mediated knockouts, DNA from edited CAR-T cells was extracted using the DNeasy Blood&Tissue kit (Qiagen, #69504), according to manufacturer’s instructions. For each locus, the region surrounding the expected cut site was amplified using forward (AAVS1: AGTTAGAACTCAGGACCAACTTATT, ZC3H12C: CAGGGGAGGAGAGTGGTGTGCT, TG: CGTGGGTGGTGAGGAGGGTGAT, ITGB8: CCCTGGCACAAAAGAAGTACTCCC, RGS2: GCCGGCTCCAGACAGTTCCATG) and reverse (AAVS1: GAATATAAGGTGGTCCCAGCTC, ZC3H12C: TACAGGGAAAGGTGGGGCCTGC, TG: CGTGGGTGGTGAGGAGGGTGAT, ITGB8: GTTCCTCCTTGGAGTGTGGCGC, RGS2: AATCCCCCTCAGGCAAGCTGCT) primers. Amplification was performed using Phusion® high-fidelity DNA polymerase (NEB, #M0530S) in an Applied Biosystems 2720 Thermal Cycler. The efficiency of knockout was assessed by Sanger sequencing and quantified by using ICE v3.0 software (Synthego) on day 10 of T-cell expansion protocol.

### Isolation of tumor infiltrating T cells (CAR-TILs)

To isolate tumor infiltrating CAR-T cells (CAR-TILs), animals were sacrificed, and subcutaneous tumors were collected and disaggregated using the Tumor Dissociation kit human (Miltenyi Biotec, #130-095-929) as per manufacturer’s protocol. Briefly, tumors were cut into 2-4 mm pieces and incubated with 100 µL of enzyme H, 12.5 µL of enzyme A and 10 µL of enzyme R in a final volume of 4.4 mL of FBS-free RPMI medium for 1 h at 37°C with continuous rotation. After incubation, cell suspensions were 100 and 40 µm filtered, washed twice with PBS and stained as described below.

Dead cell exclusion was performed using Fixable viability dye eFluor 450 (eBioscience, #65-0863-14) and murine Fc receptor (FcR) blocking reagent (Miltenyi Biotec, #130-092-575) was added to prevent nonspecific labelling. Live, CD45+ CAR-TILs were separated using FACSAriaII or FACS Aria SORP (BD) cell sorters. Sorted T cells were collected and resuspended with CAR-T medium supplemented with the same cytokines used during *in vitro* primary expansion (100 IU of IL-2, or IL-7/IL-15 for ZC3H12C CAR-TILs). CAR-TILs were rested O/N in a humidified 37 °C, 5% CO_2_ incubator. The next day, CAR-TILs were used for functional studies.

### Functional studies with CAR-T cells and CAR-TILs

1×10^5^ tumor cells were seeded in 48-well plates. For studies with isolated CAR-TILs, 1×10^4^ in 96-well plates were seeded. After overnight incubation, CAR-T cells were added at an effector-to-target ratio of 3:1 or 1:3 (effector-to-target ratio for each experiment is indicated in figure legends). At indicated experiments, anti-PD-L1 (Durvalumab) antibodies were added to CAR-TILs at a final concentration of 10 ng/mL. For cytokine secretion, supernatants were collected 24 h after co-culture and the concentration of human IFNγ, IL-2 and TNF-α was analyzed by ELISA using the following kits: DuoSet ELISA Development kit (R&D Systems, #DY285B and #DY202) and ELISAMax DeluxSet human TNF-α, following manufacturer’s protocol. Absorbance was determined using Gen5 2.07 (Biotek) or iControl 2.0 (LifeSciences) software in a microplate reader set to 450 nm. Cytokine secretion by isolated CAR-TILs after 24 h co-culture was also determined using LegendPlex Human CD8/NK Panel (Biolegend, #741187), following manufacturer’s instructions.

To analyze functionality of dysfunctional CAR-TILs after CAR-independent stimulation, cell stimulation cocktail (eBioscience, #00-4970) was added to CAR-TILs. 16 h later, supernatants were collected and cytokines were analyzed by ELISA as previously described.

Real-time cytotoxicity of CAR-T cells and CAR-TILs was analyzed using xCELLigence Real-Time Cell Analyzer System (Agilent). Briefly, 1×10^4^ tumor cells were plated in 50 µL of medium per well of an E-16 plate (Agilent, #5469830001). After 18 h, effector cells were added (effector-to-target ratio are specified in figure legends) in a final volume of 100 µL. Cell index was monitored every 20 min to up to 6 days. Relative impedance was normalized to the maximum cell index value before T cell plating. Mean cell index ±SEM of duplicates is plotted.

For intracellular staining experiments, SKOV3 tumor cells (5×10^5^) were seeded in 12-well plates and CAR-T cells were added after overnight incubation at an effector-to-target ratio of 1:3. 20h later, GolgiPlugTM (BD Bioscience, #555029) was added to each well. Cell stimulation cocktail was added to the corresponding positive control. 4 h later, flow cytometry staining was performed as described below.

### *In vitro* restimulation assay of CAR-T cells

HCC1954 tumor cells were routinely seeded in 6-well plates at 1×106 cells per well in a final volume of 4 mL of medium the day preceding T cell seeding. HER2 CAR-T cells knockout out for ZC3H12C or AAVS1, or UTD T cells were thawed and rested overnight at 4×10^6^ cells/ml in T cell medium. After overnight resting, T cells were counted by trypan blue exclusion and 3×10^6^ T cells/well were transferred in duplicates to the HCC1954 plates to final volume of 6 mL per well. After 3-4 days, the co-cultures were thoroughly resuspended by frequent pipetting and the T cell number was determined by trypan blue exclusion. Cell suspensions were collected, spun down and resuspended in fresh RPMI medium. The resulting cell suspension was transferred into HCC1954-coated plates (3×10^6^ T cells/well) for continuous co-culture. This process was repeated for 14-21 days. Each week of the restimulation assay, cytotoxicity and cytokine production by CAR-T cells was analyzed after co-culture with fresh tumors cells as explained before.

### Antibodies, flow cytometry (surface and intracellular) and cell sorting

Cell viability was determined using Fixable viability dye eFluor™ 450 (eBioscience, #65-0863-14) followed by surface antibody staining in FACS buffer. Cells were incubated with surface antibodies (**Table 1**) for 30 min at 4°C in the dark. When staining tumor samples, murine Fc receptor (FcR) were blocked using FcR blocking reagent (Miltenyi, #130-092-575) added for 10 min at 4°C prior to surface staining to prevent non-specific labelling.

Surface CAR expression was assessed using biotin-SP-AffiniPure F(ab)’2 fragment-specific goat anti-human or anti-mouse (Jackson Immunoresearch), followed by incubation with streptavidin conjugated to a fluorochrome (**Table 1**).

Intracellular staining was performed using the FoxP3/Transcription Factor Staining Buffer set (ThermoFisher, #00-5523-00), according to manufacturer’s protocol. FACS Canto 3L, Fortessa 4HT or Fortessa 5L (BD Biosciencies) were used for flow cytometry acquisition.

A FACSAria II or a FACSAria SORP (BD Biosciences) was employed for cell sorting. Samples were recovered in a 50% solution of FBS in PBS at 4°C. Flow cytometry data was analyzed using FlowJo software (v10).

### Design of a custom sgRNA library for CRISPR/Cas9-mediated knockouts

A custom library of sgRNAs targeting 300 gene candidates was purchased from Cellecta (California, United States). 1256 unique sgRNA sequences were designed, including 4 sgRNAs targeting each candidate gene and 56 control sgRNAs. Control sgRNAs included: 10 non-targeting, 10 intron-targeting (ALAS1, C1orf43, DMD, GPI and RAB7A), 10 safe harbor-targeting (AAVS1 and ROSA26) and 10 sgRNAs targeting essential genes (CDC16, GRF2B, HSPA4, HSPA9 and PAFA1HBA). As controls of positive enrichment screen, 8 sgRNAs were included targeting PD-1, LAG-3; and as controls of negative enrichment after screen, 8 sgRNAs were included targeting TCF7 and BCL-2 (4 sgRNA each). Oligonucleotide sequences were synthesized and cloned into the pRSGEG-U6-sg-EF1-TagGFP2 library vector (Cellecta). Quality controls were performed by Cellecta to confirm the configuration of the sgRNA expression cassette, ensure correct insertion and confirm full representation of the oligo pool.

### Immunohistochemistry analysis

Immunohistochemical studies we performed on formalin-fixed, paraffin-embedded tumor sections at the HCB-IDIBAPS Biobank, according to standard protocols. In brief, paraffin sections slides were developed using Immunostainer Bond Max (Leyca Biosystems). Low pH retrieval was performed (Bond ER1 Buffer solution, Leyca Biosystems) for 20 min followed by 1 h incubation at RT with primary ErbB2 (HER-2) (Invitrogen, #MA5-14509) or CD8 (Invitrogen, #MA5-14548) recombinant rabbit monoclonal antibodies. Antigen visualization was performed by incubating the sections with anti-rabbit Bond Refine Polymer (Leica Biosystems) for 10 minutes, followed by development with 3′-diaminobenzidine (DAB) as the chromogen. Nuclear counterstaining was then carried out by incubating with Harris Hematoxylin (Leica Biosystems) for 12 minutes. Images were obtained using a Nikon Eclipse E600 inverted microscope and an Olympus DP72 camera.

Quantification of CD8+ cells in IHC preparations was outsourced to the Histopathology core facility of the IRB (Barcelona). Brightfield images were acquired with a NanoZoomer-2.0 HT C9600 digital scanner (Hamamatsu) and visualized using NDP-view 2 U123888-01 software (Hamamatsu). Image analysis was performed using QuPath software v0.5.1 and a positive cell detection algorithm to maximize the detection of CD8+ cells.

### Soft clustering analysis

To study the kinetics of gene expression in CAR-TILs during preinfusion, effective and dysfunctional phases, soft clustering analysis was performed using fuzzy c-means algorithm with Mfuzz package v2.52.0 in R ^45^. In brief, gene expression data obtained by RNA sequencing was pre-processed in order to exclude genes with >25% missing measurements and variance <0.1, and to standardise gene expression to a mean of 0 and a standard deviation of 1. After pre-processing, 9.827 genes were included in the soft clustering analysis. To determine the appropriate number of clusters for our dataset, *cselection* function was used to calculate the minimum distan°ce between clusters and the partition coefficient for a number of clusters. The optimal number of clusters was finally set based on the assessment of their biological relevance.

### Bulk RNA sequencing

Total cellular RNA was isolated from effective and dysfunctional CAR-TILs and from preinfusion CAR-T cells using RNeasy Mini kit (Qiagen, #74104), as per manufacturer’s protocol. The concentration and quality of purified RNA was assessed using Qubit (Thermo Fisher). dsDNA libraries for Illumina sequencing were generated at the Genomics Unit of Center for Genomic Regulation (CRG, Barcelona) using the NEBNext® Small RNA Library Prep Set for Illumina® kit (NEB, #E7330), according to the manufacturer’s protocol. Library amplification was performed by PCR using custom Unique Dual Indexes (UDIs) and all purification steps were performed using AgenCourt AMPure XP beads (Beckman Coulter, #A63882). The quality of dsDNA libraries was analyzed using Agilent High Sensitivity DNA Kit (Agilent, #5067-4626). Size selection of DNA fragments was performed using 6% Novex TBE PAGE Gels (Thermo Fisher, #EC6265BOX) and the final pool was quantified by qPCR using the KAPA Library Quantification Kit (Kapa Biosystems, #KK4835). Libraries were sequenced on a NextSeq2000 platform with 1×50-bp paired-end reads per sample.

### Bioinformatic analysis of bulk RNA sequencing

For data analysis of RNA sequencing data, the output data from the NextSeq550, BCL files were converted into FastQ files and demultiplexes using BCL2FastQ (v2.20.0.422) using the following parameters: --barcode-mismatches 0 --no-bgzf-compression --minimum-trimmed-read-length 0 --mask-short-adapter-reads 0. The quality of sequenced reads was assessed using FastQC (v0.12.1) with default parameters. RNA-sequencing reads were mapped against GRCh38 human reference genome in combination with the sequence of the WPRE, a unique sequence in the CAR plasmid (ATCAACCTCTGGATTACAAAATTTGTGAAAGATTGACTGGTATTCTTAACTATGTTGCTCCTTTTACGCTATGTGGATACGCTGCTTTAATGCCTTTGTATCATGCTATTGCTTCCCGTATGGCTTTCATTTTCTCCTCCTTGTATAAATCCTGGTTGCTGTCTCTTTATGAGGAGTTGTGGCCCGTTGTCAGGCAACGTGGCGTGGTGTGCACTGTGTTTGCTGACGCAACCCCCAC) to identify CAR-expressing cells, using with STAR/2.7.8^46^ and using ENCODE parameters. Gene quantification was performed with HTSeq v0.11.2^47^ with the following specific parameters: --secondary-alignments ignore, --nonunique fraction.

Differential expression analysis was performed with limma v3.42.3 R package, using the voom transformation method^48^. To account for repeated measures on the same individual over time, limma’s duplicate Correlation function was used.

### Ingenuity pathway and gene-set enrichment analysis

RNA-seq data was analyzed using Ingenuity Pathway Analysis (IPA, Qiagen. Pathway analysis was performed in differentially expressed genes between dysfunctional and preinfusion or effective vs preinfusion. Upregulated genes from each comparison (p-adjusted value <0.05 and log (fold change)>1) (N=3052 for dysfunctional vs preinfusion and N=1590 for effective vs preinfusion) were sorted by p-adjusted value and the ingenuity knowledge base was used as reference set. Upstream regulator analysis was performed using upregulated genes between dysfunctional and preinfusion (N=3052). Only transcription factors previously associated with T cell exhaustion are shown.

Pre-ranked Gene-Set Enrichment Analysis (GSEA)^49^ was conducted over the ranked list of upregulated genes between dysfunctional and preinfusion phases (N=3052) based on the fold change obtained from the differential expression analysis. GSEA was performed using the fgsea package (v1.28.0) with default parameters (https://bioconductor.org/packages/release/bioc/html/fgsea.html). GSEA interrogated a chosen signature related to exhausted CD8+ T cells^50^ and the enrichment plot was generated using the same package.

### Amplicon-sequencing

To analyze enrichment of sgRNAs in T cells after screen, CD8+ CAR-T cells (expansion screen) or CAR-TILs (in vivo screen) expressing GFP (as a surrogate of library integration) were isolated and DNA was extracted using DNeasy Blood&Tissue kit (Qiagen, #69504), according to manufacturer’s protocol. As control, sgRNA representation was analyzed from the lentiviral pool by extracting lentiviral RNA using NucleoSpin RNA virus (Macherey-Nagel, #740956.10). Reverse transcription of the lentiviral RNA was performed using Transcriptor First Strand cDNA Synthesis kit (Roche, #04379012001) and a primer (AGGCAGCGCTCGCCGTGAGGA), as per manufacturer’s protocol.

DNA libraries for Amplicon-sequencing were prepared by first amplifying the insert using the high-fidelity DNA polymerase KAPA included in the KAPA HiFi HotStart ReadyMix PCR Kit (Roche, #KK2602). Forward (ACACTCTTTCCCTACACGACGCTCTTCCGATCTATTAGTACAAAATACGTGACGTAGAA) and reverse (GTGACTGGAGTTCAGACGTGTGCTCTTCCGATCTAGTAGCGTGAAGAGCAGAGAA) primers were designed to include partial P5 and P7 adapters, respectively, for subsequent Illumina sequencing. For each sample, between 270 and 1000 ng of genomic DNA were amplified by preparing from 3 to 10 PCR reactions with up to 100 ng of genomic DNA per reaction to maximize fidelity during amplification. Amplification was performed in an Applied Biosystems 2720 Thermal Cycler and PCR products were confirmed by electrophoreses in agarose gel. Quality and quantity of DNA samples were assessed using NanoDrop 1000 spectrophotometer (Thermo Fisher).

Second PCR was performed using NEBNext Q5 Hot Start HiFi PCR Master Mix (NEB, #M0543L) and index 2 (i5) primers containing Illumina’s P5 and P7 sequences. Final libraries were analyzed using Agilent Bioanalyzer High Sensitivity assay (Agilent, #5067-4626) to estimate the quantity and check size distribution, and were then quantified the KAPA Library Quantification Kit KK4835 (Roche, #07960204001). Libraries were prepared at the Center for Genomic Regulation (CRG, Barcelona) using standard protocols. DNA libraries were sequenced 1 * 51+10+10 bp on Illumina’s NextSeq500 using a custom sequencing primer (TCTTGGCTTTATATATCTTGTGGAAAGGACGAAACACCG) for read 1 with a threshold of 13 million reads.

### MAGeCK analysis

Amplicon-seq data were analyzed using the MAGeCK algorithm (v.0.5.9.5)^51^ via the Singularity image built using the command singularity build mageck.sif. The MAGeCK algorithm was used to perform normalization and to calculate log fold changes of genes and sgRNAs, positive and negative gene rankings and P values. Count normalization was performed by total counts and gene log fold changes were calculated by using second best method.

For the quality control analysis, one sample from plasmid DNA was compared to one sample of lentiviral DNA and with five T cell samples collected on day 4 of *in vitro* primary expansion before Cas9-mediated knockout (five different healthy donors). For the expansion screen, five T cell samples were collected on day 4 of *in vitro* primary expansion before Cas9 electroporation and compared to four T cell samples collected on day 10-11 after primary activation. For the in vivo screen, four CAR-TIL samples were isolated from tumors on day 20-28 (3 different healthy donors) and compared to T cells before Cas9-mediated electroporation. Paired analysis by donor was performed for the *in vivo* screen.

### Single-cell RNA and ATAC-sequencing

A minimum of 100.000 tumor infiltrating CD8+ T cells were isolated at effective and dysfunctional phase or harvested prior to infusion (preinfusion) using the *in vivo* model of CAR-T cell dysfunction and the isolation protocol described in “Isolation of CAR-TILs” section.

Nuclei isolation was performed at CNAG-CRG (Barcelona) following the Nuclei Isolation for Single Cell Multiome ATAC + Gene Expression Sequencing protocol (10X Genomics; CG000365). After isolation, nuclei were resuspended at a concentration of 925-2,300 nuclei/μL and counted by trypan blue exclusion. Single cell RNA and ATAC-seq libraries were prepared following the Chromium Next GEM Single Cell Multiome ATAC + Gene Expression User Guide (10X Genomics; CG000338). Transposed nuclei were partitioned into Gel Bead-In-Emulsions (GEMs) by using the Chromium Controller with Chip J. Amplified cDNA was quantified on an Agilent Bioanalyzer High Sensitivity chip (Agilent Technologies) and used for both single cell RNA and ATAC seq library generation. Single cell libraries were indexed with 13 cycles of amplification using the Dual Index Plate TT Set A (10X Genomics; PN-3000431). ATAC-seq libraries were indexed with 8 cycles of amplification using the Sample Index N Set A (10X Genomics; PN 3000427). Size distribution and concentration of full-length libraries were verified on an Agilent Bioanalyzer High Sensitivity chip (Agilent Technologies).

Sequencing was carried out on a NovaSeq 6000 sequencer (Illumina) using the following settings: 28 bp (Read 1) + 8 bp (i7 index) + 0 bp (i5 index) + 89 bp (Read 2) for single cell RNA, and 50 bp (Read 1N) + 8 bp (i7 Index) + 16 bp (i5 Index) + 49 bp (Read 2N) for ATAC-seq, to obtain approximately 40,000 paired-end reads per cell.

### Bioinformatic analysis of single-cell RNA seq and ATAC-seq

Processing of single cell multiome data was performed using 10X Genomics Cell Ranger ARC 2.0.2. Sequencing reads were aligned to the human reference genome (ARC multiome GRCh38 2020-A-2.0.0, 10x Genomics). To obtain a valid set of cells for analysis, cells were filtered to exclude potential doublets using DoubletFinder, minimum of 500 counts and 500 genes per cell, maximum of 10% of mitochondrial genes counts, and a novelty score higher than 0.8. Following primary processing, beween 6.000 and 12.000 cells per sample were retained.

Differential expression analysis were performed in R (version 4.5.1) using the Seurat (version 4.3). Read counts were normalized using the SCTransform method and processed downstream for Uniform Manifold Approximation and Projection (UMAP). Clusters were identified by multimodal neighbors using the WNN method. DNA accessibility was analyzed using Signac (v1.14). For that, a peak calling was performed, a unified set of peaks were generated, filtered (peak widths > 20 and < 1000) and quantified.

To annotate the cell types of CAR-TILs samples, the scRNA-seq datasets were mapped on gene signatures reported by Zheng and colleagues^25^. Scalar transformation was applied to the expression values of each gene across all cells within each gene set. For each cell, the transformed values were subsequently summed to generate a gene set-specific score. Cell states were assigned based on a percentile threshold: cells with scores exceeding the 75th percentile for a given gene set were annotated with the corresponding cell type. In cases where a cell surpassed the threshold for multiple gene sets, the cell type associated with the highest score was selected.

Motif analysis was performed in cells ZC3H12C-positive cells, defined as those with normalized count values above the 50^th^ percentile. A set of peaks was first identified based on either their unique presence in ZC3H12C-positive cells or a log2 fold change >1 when compared to ZC3H12C-negative cells. Differentially expressed genes were then identified based on a log2 fold change > 0.13 and a p-value < 0.05 from the same comparison. A subset of peaks located nearest to these differentially expressed genes was selected for motif analysis. Motif enrichment analysis was conducted using the JASPAR2020 database. Motifs were considered significantly enriched if they exhibited a fold enrichment > 1.5 and an adjusted p-value < 0.001, relative to a background set of randomized genomic regions matched for size and GC content.

### Statistical analysis

All statistical analyses were performed using GraphPad Prism v10.4.1 (GraphPad software Inc.). For comparisons of two groups, two-tailed paired or unpaired tests were used. One-way analysis of variance (ANOVA) with Tukey’s multiple comparisons test was used for comparing three or more groups in a single condition. For the analysis of three or more groups in multiple conditions, a two-way ANOVA with Sidak or Tukey correction were performed. Asterisks indicate statistical significance as follows: ns, not significant; *p<0.05; **p<0.01; ***p<0.001; ****p<0.0001.

## Data availability

Bulk RNA-sequencing data is deposited at the GEO database with accession number GSE309402. Amplicon-sequencing and single-cell RNA sequencing are pending to upload.

## Acknowledgements

This work received funding from the Spanish Ministry of Science and Innovation under a Ramon y Cajal grant (RYC2018-024442-I to S.G.) and Retos Investigación (PID2019-109546RA-I00-PI), the Innovative Medicines Initiative 2 Joint Undertaking under grant agreement No 116026 (this Joint Undertaking receives support from the European Union’s Horizon 2020 research and innovation program and EFPIA), the Horizon European Innovation Council 2021 Pathfinder Challenges-01 under grant agreement No 101070740, “la Caixa” Foundation under the grant agreement LCF/PR/SP23/52950004, the Spanish Association Against Cancer (LABAE20022GUED to S.G., INVES222988 RODR to A.R-G.). A.P. received funding from Fundación CRIS contra el cáncer PR_EX_2021-14, Agència de Gestó d’Ajuts Universitaris i de Recerca 2021 SGR 01156, Fundación Fero BECAONCOXXI21, Instituto de Salud Carlos III PI22/01017, Asociación Cáncer de Mama Metastásico IV Premios M. Chiara Giorgetti, Breast Cancer Research Foundation BCRF-23-198, and RESCUER, funded by European Union’s Horizon 2020 Research and Innovation Programme under Grant Agreement No.847912. P.B. received a “Formación de Personal Universitario” grant to conduct predoctoral studies (FPU19/01931).

This study was developed at Institut d’Investigacions Biomèdiques August Pi i Sunyer (IDIBAPS). We are indebted to the Biobank and to the Flow Cytometry and Cell Sorting core facilities at IDIBAPS and to the animal facility of the Universitat de Barcelona for their technical help. We thank the Genomics Unit at the CRG for assistance with the sequencing. We thank Holger Heyn and Domenica Marchese for their help with the Multiome. We thank the Functional Genomics team at CNAG for their technical help during the analysis of RNA seq data. We thank Julia Ponomarenko for the bioinformatic analysis with MAGeCK. We thank the Histolopathology core facility at IRB for their technical help with the quantification of immunohistochemistry preparations. Institutional support to CNAG was provided by the Spanish Ministry of Science and Innovation through the Instituto de Salud Carlos III, and by the Generalitat de Catalunya through the Departament de Salut and the Departament de Recerca i Universitats.

## Authors contribution

P.B. designed and performed experiments, analyzed and interpreted the data, and wrote the manuscript. A.R-G. designed and performed experiments. A.G-A. analyzed single cell seq and ATAC seq data. M.G-A., P.C., M.B., G.C., I.A-S. and M.S-C. provided technical assistance with *in vitro* experiments. T.L-J. performed bioinformatic analysis. G.C. and J.C. performed *in vivo* experiments. B.M. developed some of the CAR constructs used in the study. A.E-C. analyzed bulk RNA seq and supervised the soft clustering analysis. A.P., A.U-I. and L-G. provided conceptual guidance. M.M-P. provided conceptual guidance on bioinformatic analysis. S.G. supervised the project, including design of experiments, data analysis, and manuscript writing. All authors revised the manuscript.

