## Supplementary data for "*In vivo* CRISPR-based screen identifies ZC3H12C as a mediator of CAR-T cell dysfunction in solid tumors"

**Supplementary material**

**FIGURE S1**

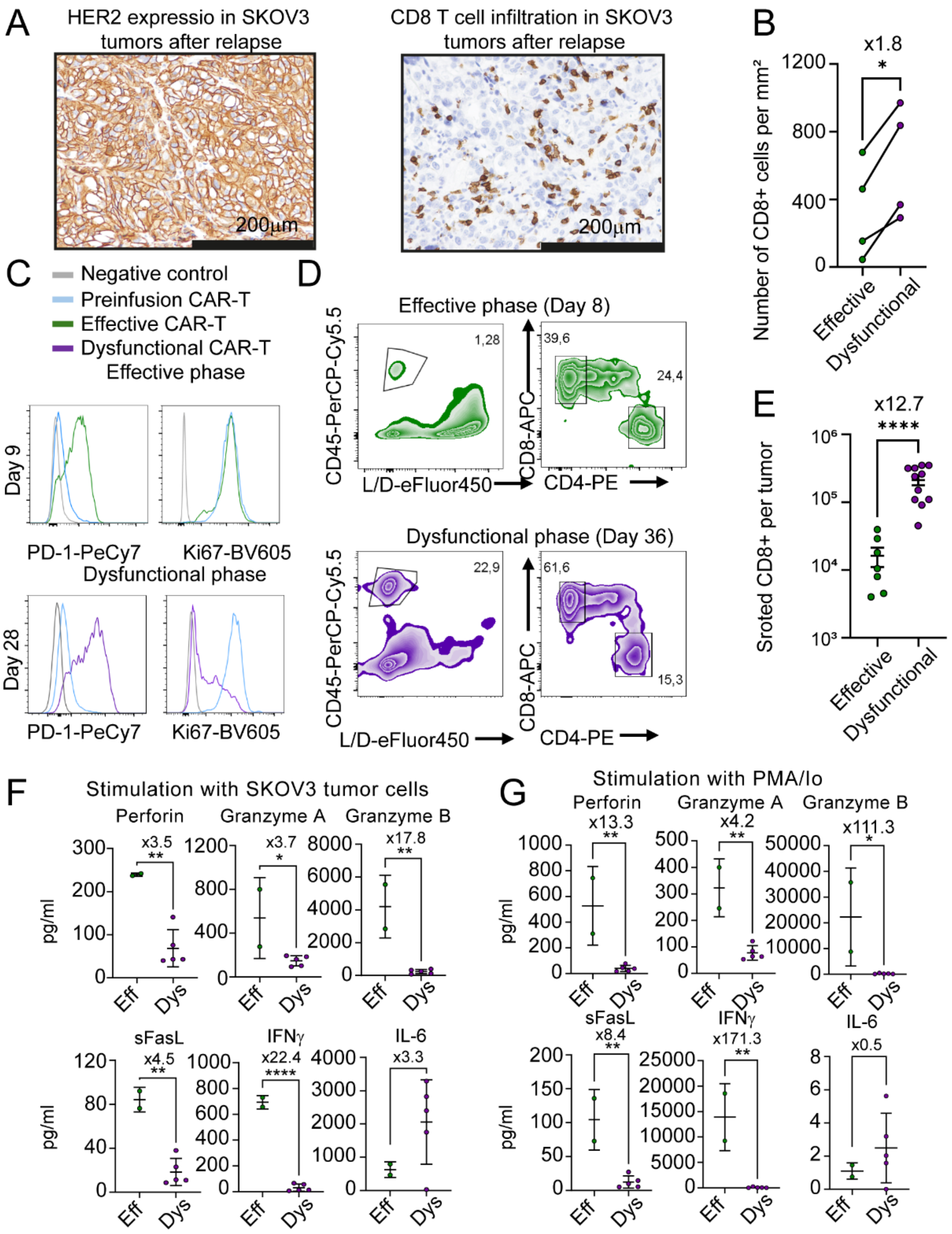


**Figure S1. HER2-CD28z CAR-T cells accumulate in ovarian xenografts overtime.**

**A**) Immunohistochemical (IHC) staining of CD8 or HER2 in SKOV3 tumors that escape to CAR-T cell therapy. Scale bar, 200µm. Representative images from n=4 in IHC analysis. **B**) Number of CD8+ cells per tumor area as quantified from brightfield images of immunohistochemical staining in tumors treated with HER2-28z CAR-T cells. Each dot represents a different healthy donor (n=4 paired donors). *p<0.05 by two-tailed paired t-test. Fold change relative to effective phase is indicated. **C**) Flow cytometry analysis of PD-1 and Ki67 expression in effective compared to dysfunctional CAR-TILs and to preinfusion product. Graphs of a representative experiment. **D**) Flow cytometry analysis of infiltrating T cells at effective and dysfunctional phase in SKOV3 tumors treated with HER2-28z CAR-T cells. **E**) Number of CD8+ T cells (gated on live, CD45+) isolated from SKOV3 tumors at effective and dysfunctional phase after treatment with CAR-T cells by FACS-sorting. Each dot represents an individual tumor. Data are plotted as mean ±SEM sorted cells per tumor (n=5 healthy donors). ***p<0.001 by two-tailed unpaired t-test. Fold change relative to dysfunctional is indicated. **F-G**) Cytokine release by effective (EFF) and dysfunctional (DYS) CAR-TILs following *ex vivo* SKOV3 co-culture (E:T=3:1) (F) or stimulation with PMA/Ionomycin (G) by HER2-28z CAR-TILs isolated from SKOV3 tumors at effective and dysfunctional phase after therapy. Data are plotted as mean ±SD of 2 healthy donors. *p<0.05, **p<0.01 by two-tailed unpaired t-test. Fold change relative to effective is indicated.

### FIGURE S2

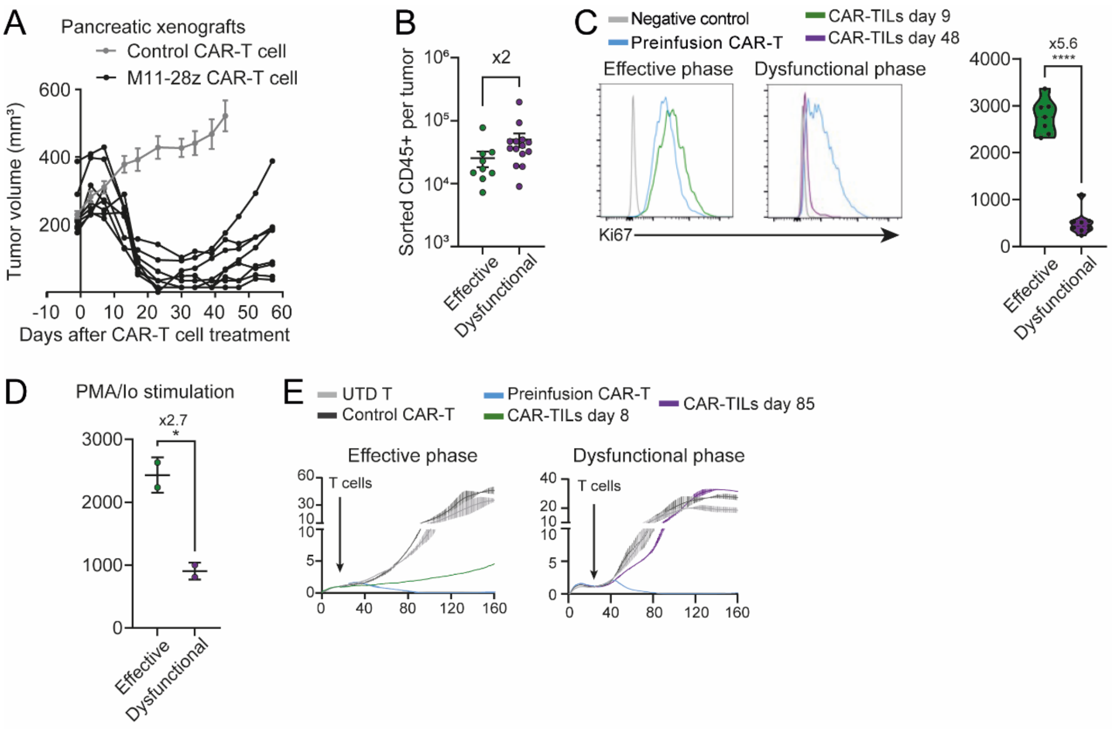


**Figure S2. M11-28z CAR-T cells become dysfunctional in an *in vivo* model of pancreatic xenografts causing tumor escape.**

**A**) Tumor measurements of NSG mice bearing subcutaneous CAPAN-2 xenograft tumors treated with $2x10^6$ M11-28z CAR-positive T cells or CD19-28z CAR-positive T cells as a control. Data are plotted as mean ±SEM tumor volume or as individual tumors (n=6-8 tumors per group). **B**) Number of CD45+ cells isolated from SKOV3 tumors at effective and dysfunctional phase by FACS-sorting. Data are plotted as mean ±SEM of sorted cells per tumor (n=2 healthy donors). Fold change in effective relative to dysfunctional is indicated. **C**) Flow cytometry analysis of Ki67 expression in effective compared to dysfunctional CD8+ M11-28z CAR-TILs and to preinfusion product (gated in live, CD45+). Graphs of a representative experiment (left). Violin plot showing mean fluorescence intensity of Ki67 in effective compared to dysfunctional CAR-TILs (right). Each dot represents a tumor (n=4 healthy donors). ****p<0.0001 by two-tailed unpaired paired t-test. Fold change relative to dysfunctional phase is indicated. **D**) IFNγ production by M11-28z CAR-TILs after *ex vivo* stimulation with PMA/Ionomycin. Data are plotted as mean ±SD in 2 different healthy donors. *p<0.05 by two-tailed paired t-test. **E**) *Ex vivo* real-time cytotoxicity analysis of isolated M11-28z CAR-TILs at effective (left) and dysfunctional (right) phase against CAPAN-2 tumor cells (E:T=2:1). Data are plotted as mean ±SEM normalized cell index of a representative experiment. Time when T cells were added is indicated.

**FIGURE S3**

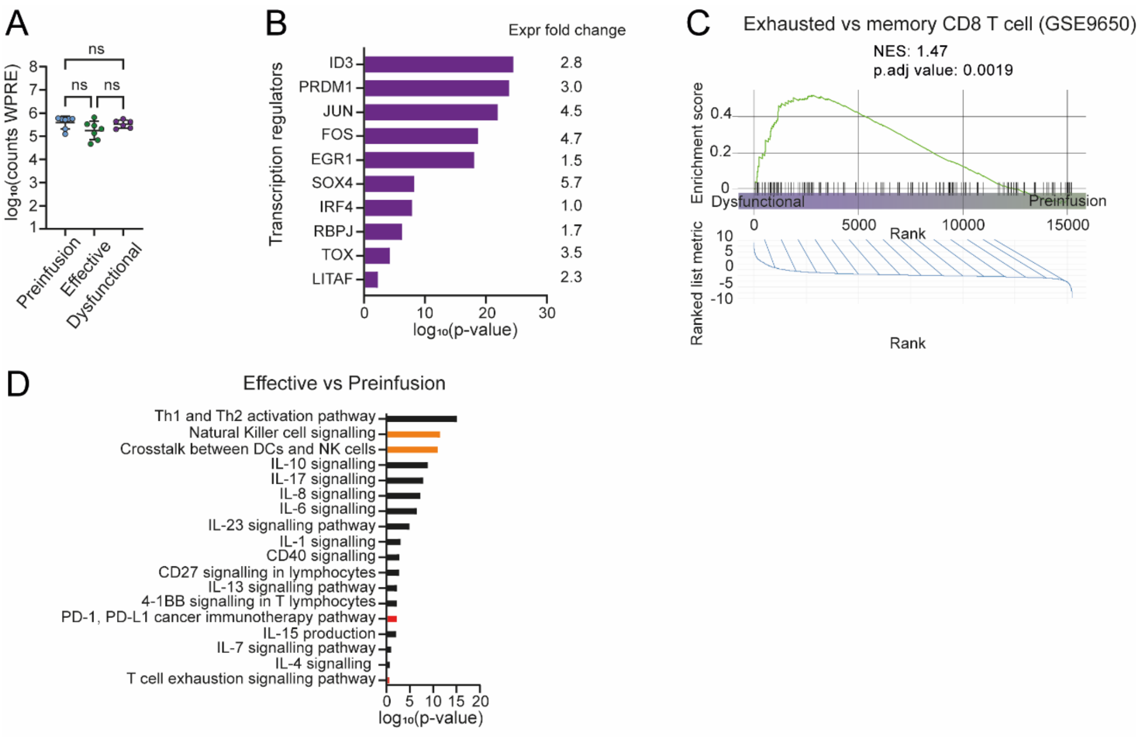


**Figure S3. CAR-T cells upregulate signatures of T cell exhaustion after chronic antigen exposure *in vivo*.**

**A)** Analysis of WPRE counts (present in CAR mRNA) from RNA-seq data in effective compared to dysfunctional CAR-TILs and to preinfusion product. Each dot represents a sample. n.s. non-significant by one-way ANOVA. **B**) IPA Upstream Regulator Analysis of transcription factors predicted to regulate the genetic profile in dysfunctional CAR-TILs. Selected transcription factors related to exhaustion are ranked by p-value. Fold change of expression from the differential expression analysis of dysfunctional versus preinfusion phase is shown. **C**) Gene Set Enrichment Analysis (GSEA) of upregulated genes in exhausted CD8 T cells from **Wherry et al. Immunity. 2007** in dysfunctional versus preinfusion CAR-TILs. NES: normalized enrichment score, P: nominal p-value given by the GSEA program. **D**) IPA analysis of upregulated genes in effective compared to preinfusion samples. Selected pathways are ranked by p-value of enrichment. Orange bars denote NK cell-related pathways, and red bars indicate pathways of T cell exhaustion.

### FIGURE S4

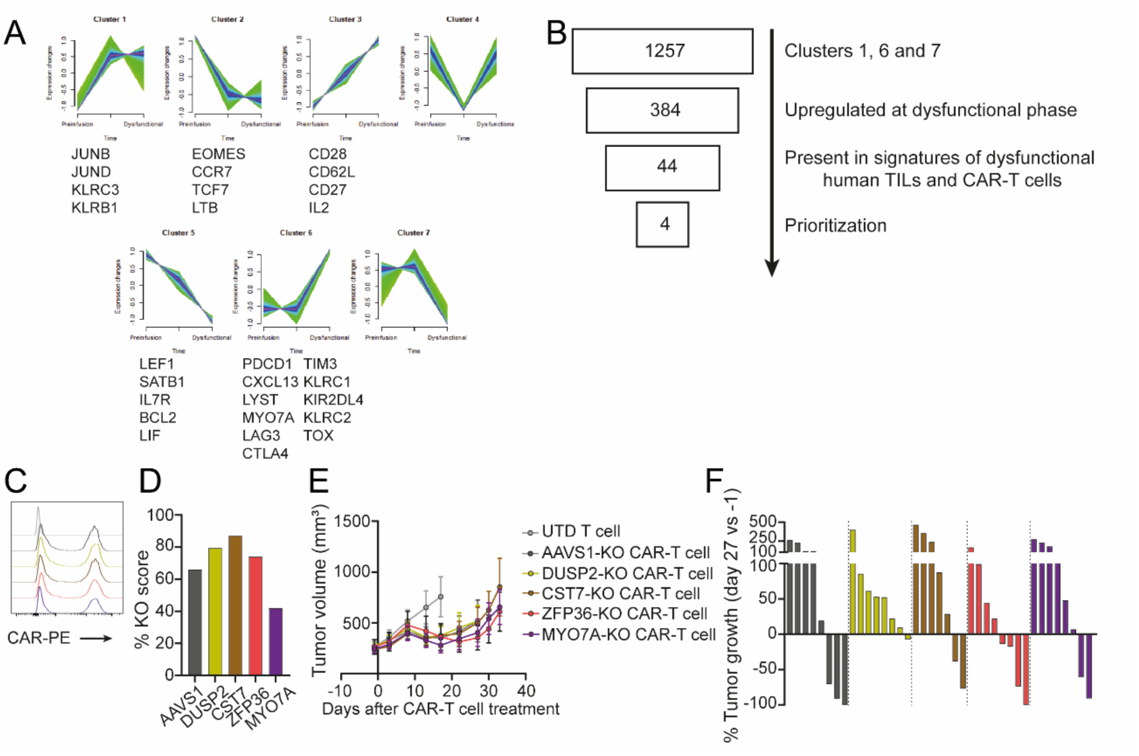


**Figure S4. *In vivo* evaluation of candidate genes in HER2-28z CAR-T cells identified through literature search.**

**A**) Soft clustering analysis of RNAseq data on preinfusion, effective and dysfunctional CAR-TILs. NK cell markers and relevant genes associated with T cell exhaustion, memory and proliferation belonging to each cluster are indicated. **B**) Pipeline for the selection of candidate genes based on existing literature. **C-F**) NSG mice bearing SKOV3 tumors were infused with a single dose of 2x10$^{6}$ CAR-positive T cells with a knockout in DUSP2, CST7, ZFP36 or MYO7A. AAVS1-KO CAR-T and UTD T cells were used as control. **C**) Histograms of CAR expression as analyzed by flow cytometry. **D**) KO efficiency was analyzed using ICE software (Synthego) and represented as KO score. **E**) Tumors were measured at indicated timepoints. Data are plotted as mean ±SEM of tumor volume (n=1 healthy donor, 8-10 tumors per group). **F**) Change in tumor volume on day 27 compared to baseline in mice for each group is shown.

**FIGURE S5**

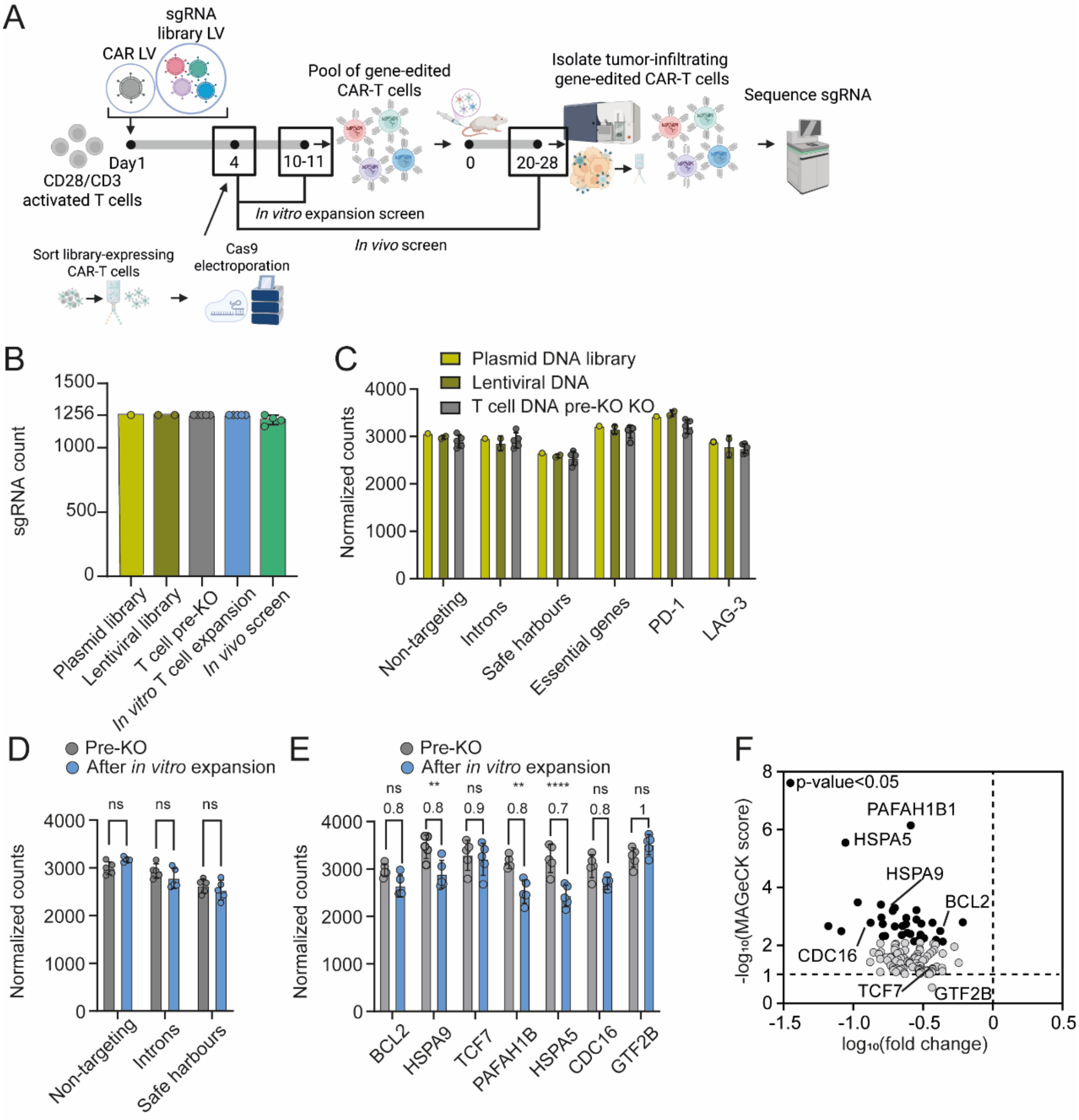


G

| Gene | Ranking |
|---|---|
| ZC3H12C | 1 |
| UCP3 | 2 |
| AHNAK2 | 3 |
| STON1 | 4 |
| MYH6 | 5 |
| LILRB1 | 6 |
| PTK7 | 7 |
| TG | 8 |
| INSRR | 9 |
| ADAM12 | 10 |
| COL4A1 | 11 |
| ITGB8 | 12 |
| FN1 | 13 |
| ACTG2 | 14 |

**Figure S5. Validation of the *in vivo* CRISPR screen strategy to identify mediators of CAR-T cell dysfunction.**

**A**) Schematic of experimental procedure to generate and test a pool of KO-CAR-T cells. Squares indicate which samples were compared to analyze sgRNA enrichment during *in vitro* primary expansion of pooled KO CAR-T cells or after *in vivo* screen. **B**) Number of sgRNAs as determined by Amplicon-sequencing of plasmid DNA library, pooled lentiviral DNA, T cells before Cas9-mediated KO, pooled KO CAR-T cells after *in vitro* expansion and pooled-KO CAR-T cells after *in vivo* screen. Data is plotted as mean ±SEM of sgRNA count. **C**) Abundance of negative control sgRNAs (non-targeting, intron-targeting and safe harbour) and positive control sgRNAs (essential genes, PD-1 and LAG-3) was analyzed in the plasmid DNA, the lentiviral pooled product and in T cells transduced with the library and before Cas9 nucleofection from Amplicon-Seq. Data shows mean ±SEM of normalized sgRNA counts in each sample. **B** and **C**) Each dot represents a sample (n=1 for plasmid library, n=2 samples for lentiviral DNA, n=5 healthy donors for T cell samples after *in vitro* expansion and n=3 healthy donors for T cells after *in vivo* screen). **D-E**) Normalized abundance of control sgRNAs (**D**) or sgRNAs targeting essential genes (**E**) in T cells transduced with the library before Cas9 electroporation compared to pooled knockout CAR-T cells after *in vitro* primary expansion. Data are plotted as mean ±SEM of normalized counts. Each dot represents a sample (n=5 healthy donors). n.s. non-significative, **p<0.01, ****p<0.0001 by two-way ANOVA with Sidák's multiple comparisons test. Fold change of edited compared to non-edited CAR-T cells is indicated. **F**) Correlation between MAGeCK score and fold change from MAGeCK analysis of depleted genes in pooled KO-CAR-T cells after *in vitro* screen. Black dots indicate significantly depleted genes. **G**) Ranking of significantly enriched genes after *in vivo* screen using MAGeCK software.

**FIGURE S6**

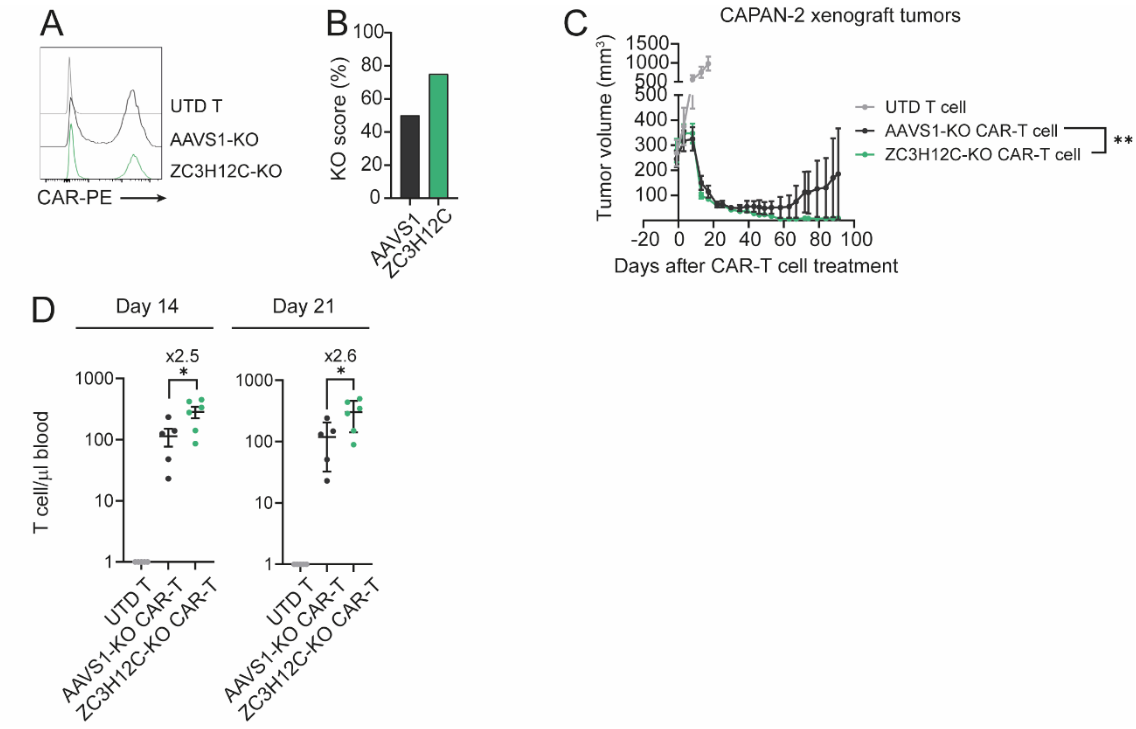


**Figure S6. ZC3H12C-KO Meso-28z CAR-T cells eliminate pancreatic xenografts and increase T cell persistence *in vivo.***

**A-D**) NSG mice bearing pre-established CAPAN-2 tumors were treated with a single dose of $2x10^{6}$ M11-28z CAR-positive T cells with genetic ablation of ZC3H12C, UCP3 or for the safe harbor AAVS1 as control. **A**) Histogram of CAR expression in M11-28z CAR-T cells used for *in vivo* experiments. **B**) Efficiency of CRISPR/Cas9-mediated knockout in M11-28z CAR-T cells as quantified using ICE tool (Synthego). **C**) Tumor volume of mice treated with KO CAR-T cells was analyzed at indicated timepoints. Data are plotted as mean ±SEM tumor volume (n=1 healthy donors, 4-5 tumors per group). *$p<0.05$, **$p<0.01$ by two-way ANOVA with Tukey's multiple comparisons test at day 91 after CAR-T therapy. **D**) Total T cell concentration in the blood of animals treated with M11-28z CAR-T cells knocked out for candidate genes or AAVS1 as control was analyzed at day 14 (left) and 21 (right) after CAR-T therapy. Each dot represents a mouse. Data are plotted as mean ±SD number of T cells (n=1 healthy donor, 4-5 mice per group). *$p<0.05$ by one-way ANOVA with Tukey's multiple comparisons test. Fold change relative to AAVS1 group is indicated.

### FIGURE S7

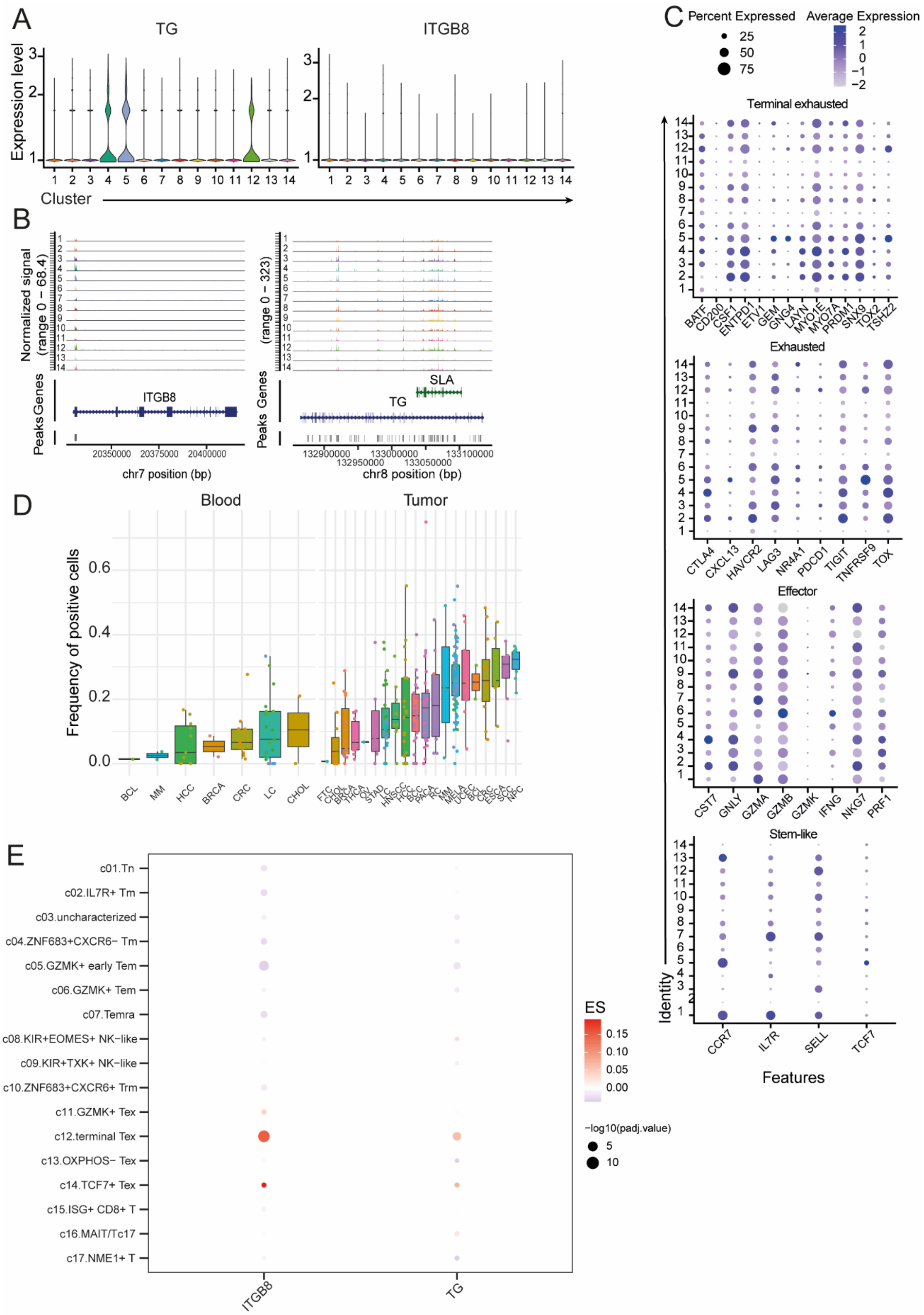
A
TG
ITGB8
Expression level
Cluster
B
Normalized signal (range 0 – 68.4)
(range 0 – 323)
Peaks Genes
ITGB8
TG
SLA
chr7 position (bp)
chr8 position (bp)
C
Percent Expressed
25
50
75
Average Expression
Terminal exhausted
BATF
CD200
CSF1
ENTPD1
ETV1
GEM
GNG4
LAYN
MYO1E
MYO7A
PRDM1
SNX9
TOX2
TSHZ2
Exhausted
CTLA4
CXCL13
HAVCR2
LAG3
NR4A1
PDCD1
TIGIT
TNFRSF9
TOX
Effector
CST7
GNLY
GZMA
GZMB
GZMK
IFNG
NKG7
PRF1
Stem-like
CCR7
IL7R
SELL
TCF7
Identity
Features
D
Blood
Tumor
Frequency of positive cells
BCL
MM
HCC
BRCA
CRC
LC
CHOL
E
c01.Tn
c02.IL7R+ Tm
c03.uncharacterized
c04.ZNF683+CXCR6− Tm
c05.GZMK+ early Tem
c06.GZMK+ Tem
c07.Temra
c08.KIR+EOMES+ NK−like
c09.KIR+TXK+ NK−like
c10.ZNF683+CXCR6+ Trm
c11.GZMK+ Tex
c12.terminal Tex
c13.OXPHOS− Tex
c14.TCF7+ Tex
c15.ISG+ CD8+ T
c16.MAIT/Tc17
c17.NME1+ T
ITGB8
TG
ES
−log10(padj.value)

**Figure S7. Single cell RNA and ATAC sequencing analysis of preinfusion CAR-T cells and of CAR-TILs isolated at effective and dysfunctional phase.**

**A**) Violin plots showing the expression of TG and ITGB8 in clusters identified by single cell RNA seq analysis. **B**) Representation of ATAC-seq tracks at TG and ITGB8 regions for cells in each cluster shown in A. **C**) Dot plots showing expression of signature markers reported by Zheng and colleagues (**Zheng, et al. Science. 2021**) in each cluster identified by single cell RNAseq. The percentage of expression is calculated as the number of cells positive for each marker divided by the total number of cells in each time point (preinfusion, effective or dysfunctional phase). The average expression is color-coded. **D-E**) Expression of candidate genes in blood and tumor from cancer patients in the pan-cancer atlas published by Zheng and colleagues (**Zheng, et al. Science. 2021**). **D**) Frequency of positive cells for ZC3H12C in blood and tumor from cancer patients. **E**) Dot plot showing expression of candidate genes in meta-clusters 1-17 the pan-cancer T cell atlas by Zheng and colleagues. Color indicates effect size (ES) and size indicates statistical significance.

**FIGURE S8**

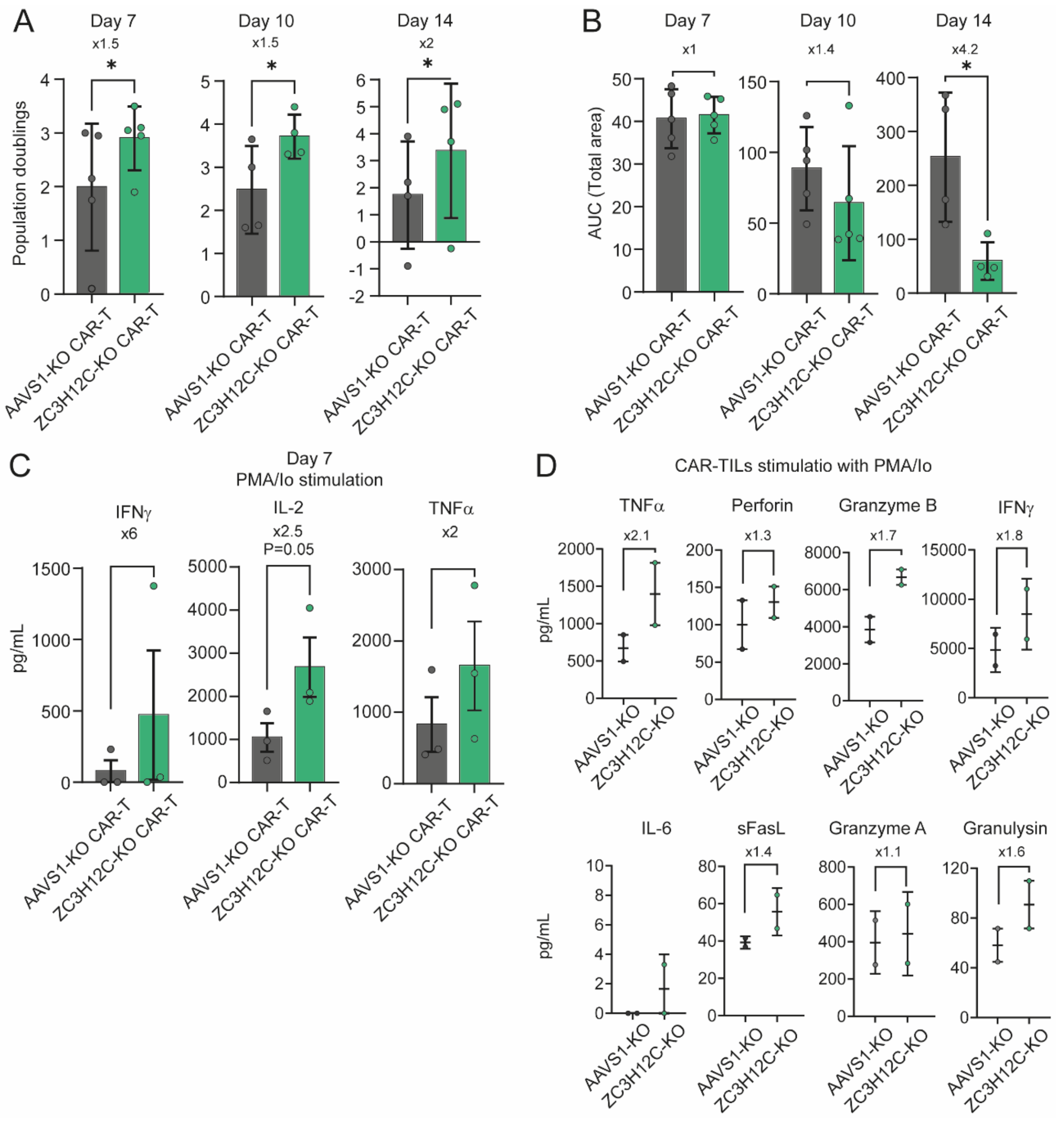


**Figure S8. ZC3H12C-KO HER2-28z CAR-T cells show superior proliferation and cytokine production after chronic antigen stimulation.**

**A-C**) AAVS1-KO and ZC3H12C-KO HER2-28z CAR-T cells were *in vitro* co-cultured with HCC1954 tumor cells (E:T=3:1). Every 3-4 days, T cells were counted and plated with fresh tumor cells. **A**) Proliferation of AAVS1-KO and ZC3H12C-KO CAR-TILs during a restimulation assay. Data are plotted as mean ±SD population doublings at indicated days of the restimulation assay (n=5 in days 7 and 10; n=4 in day 14 healthy donors). *p<0.05 by two-tailed paired t-test. Fold change relative to AAVS1 is indicated. **B**) Cytotoxicity of restimulated CAR-T cells was analyzed using a real-time assay. For each timepoint, the total area under the curve (AUC) is represented as absolute numbers. Data are plotted as mean

±SD. Each dot represents a healthy donor (n=5 for day 0 and day 7; n=4 for day 14). *$p<0.05$ by two-tailed paired t-test. **C**) Cytokine release by CAR-T cells at 7 days of stimulation with HCC1954 tumor cells following PMA/Ionomycin stimulation. Data are plotted as mean ±SD. Each dot represents a healthy donor (n=2). Fold change relative ZC3H12C-KO is indicated. **D**) CAR-TILs were isolated from SKOV3 tumors of mice after two weeks of treatment with AAVS1-KO or ZC3H12C-KO HER2-CD28 CAR-T cells. After overnight resting, CAR-TILs were stimulated with PMA/Io and cytokines were analyzed in the supernatant. Data are plotted as absolute numbers ±SD (n=2 healthy donors). Fold change relative to ZC3H12C-KO group is indicated.
